# Combined model-free and model-sensitive reinforcement learning in non-human primates

**DOI:** 10.1101/836007

**Authors:** Bruno Miranda, W. M. Nishantha Malalasekera, Timothy E Behrens, Peter Dayan, Steven W. Kennerley

## Abstract

Contemporary reinforcement learning (RL) theory suggests that potential choices can be evaluated by strategies that may or may not be sensitive to the computational structure of tasks. A paradigmatic model-free (MF) strategy simply repeats actions that have been rewarded in the past; by contrast, model-sensitive (MS) strategies exploit richer information associated with knowledge of task dynamics. MF and MS strategies should typically be combined, because they have complementary statistical and computational strengths; however, this tradeoff between MF/MS RL has mostly only been demonstrated in humans, often with only modest numbers of trials. We trained rhesus monkeys to perform a two-stage decision task designed to elicit and discriminate the use of MF and MS methods. A descriptive analysis of choice behaviour revealed directly that the structure of the task (of MS importance) and the reward history (of MF and MS importance) significantly influenced both choice and response vigour. A detailed, trial-by-trial computational analysis confirmed that choices were made according to a combination of strategies, with a dominant influence of a particular form of model sensitivity that persisted over weeks of testing. The residuals from this model necessitated development of a new combined RL model which incorporates a particular credit assignment weighting procedure. Finally, response vigor exhibited a subtly different collection of MF and MS influences. These results provide new illumination onto RL behavioural processes in non-human primates.

## Introduction

Reinforcement learning (RL) is a theoretical framework for how agents interact with their environment, predicting and optimizing summed rewards over an extended future (1). In such contexts, learned models of the environment, like Tolman’s cognitive map (2), characterize the structure of the task: for instance, reporting how actions determine both rewards and (probabilistic) changes in the state of the world. RL encompasses many methods for learning and planning. Model-free (MF) approaches learn estimates of the long-run summed reward, often by enforcing a form of self-consistency along observed trajectories between actions and subsequent states (i.e., samples reflecting the state-transition structure). MF-RL typically requires substantial sampling from the world to achieve good performance, and is therefore, like behavioural habits (3), slow to adapt to environmental change. Pure model-based (MB) approaches use the model to plan, for instance by simulating possible trajectories. Their estimates of long-run rewards are thereby readily adaptive to environmental change, just like goal-directed actions (4, 5).

MF and MB RL occupy opposite points on the spectrum of computational simplicity and statistical efficiency (6, 7). This originally inspired ideas that their output should be combined (8). Recently, rather complex patterns of interaction have been investigated, including MB training of MF (9, 10), MF control over MB calculations (11–13), the incorporation of MF values into MB calculations (14) and, of particular relevance for the present study, the creation of sophisticated, model-dependent, representations of the task that enable MF methods to work more efficiently (15), and potentially less susceptible to distraction (16). We deem these various interactions model-sensitive (MS), saving model-based for the original notion of prospective planning. Since we focus only on behavioural data, we do not attempt to unpick the particular forms of model-sensitivity that our subjects exhibit; we regard as MS any dependencies that are associated with the structure of the task rather than purely previous rewards.

Traditional studies of MF and MS strategies in rodents exploited manipulations such as outcome devaluation (17). However, these offer only limited opportunities to explore continuing tradeoffs. More recently, a class of new tasks has been invented for human subjects (8, 10) that use a state-transition structure in combination with changing outcomes to examine how the strategies are combined. However, inevitable limitations in the length of these experiments leaves us uncertain: about the stability and goodness of fit of such combinations in the long run (16); about possible implications for relatively noisy output measures, such as reaction times, which can reflect the tradeoff between speed and accuracy that separate various strategies; about the wider spectrum of MS methods; about additional facets that are routinely added to MF and MS accounts in order to fit behavioral data well, such as a bias towards preseveration; and indeed about generalization to other species. Further, characteristically different forms of MF learning have been found in primates (18) and rodents (19), motivating further investigation. Here, two rhesus monkeys were trained to perform a two-stage decision task (Fig. 1; see *SI Methods* for details) intended to induce trial-by-trial adjustments in choice that combine aspects of MF and MS learning. We used RL-based methods to analyse quantitatively several orders of magnitude more behavioural data than previous human studies, and found sensitivity to reward history (of MF and MS importance) as well as information about the state-transition structure (of MS relevance). Both forms of RL were persistently influential in the long run, and also both influenced the alacrity of responding, in agreement with the speed-accuracy trade-off associated with their computations. Our results enrich modern views of MF and MS integration (6, 20–22).

**Fig. 1.**
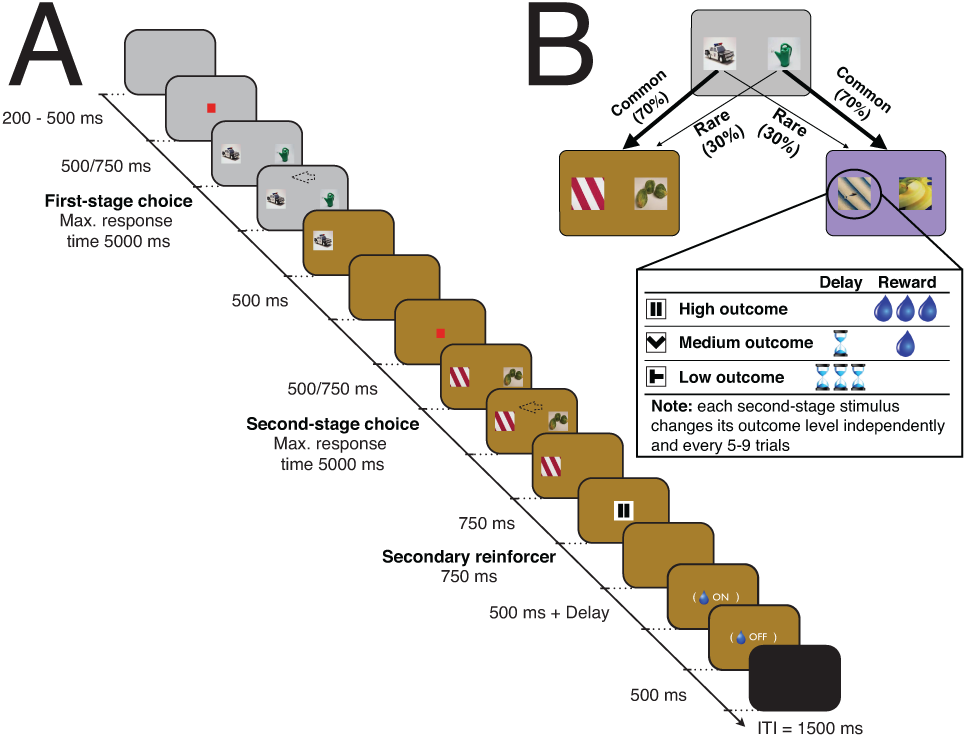
Two-stage decision task. (A) Timeline of events. Eye fixation was required while a red fixation cue was shown, otherwise subjects could saccade freely and indicate their decision (arrow as an example) by moving a manual joystick in the direction of the chosen stimulus. Once the second-stage choice had been made, the nature of the outcome was revealed by a secondary reinforcer cue (here, the pause symbol represents high reward). Once the latter cue was off the screen, there was a fixed 500 ms delay and the possibility of a further delay (for both medium and low rewards) before juice was provided (for both high and medium rewards). (B) The state-transition structure (kept fixed throughout the experiment). Each second-stage stimuli had an independent reward structure: the outcome level (defined by the magnitude of the reward and the delay to its delivery) remained the same for a minimum number of trials (a uniformly distributed pseudorandom integer between 5 and 9) and then, either stayed in the same level (with one-third probability) or changed randomly to one of the other two possible outcome levels.

## Results

The subjects performed a two-stage decision task (subject C: 15585 trials over 30 sessions; subject J: 14664 over 27 sessions), similar to the one used in a previous human study (8). In brief, two decisions had to be made on each trial (Fig. 1). At the first-stage state (represented by a grey background), the choice was between two options presented as stimuli (fixed throughout the entire task). The consequence was a transition to one of two second-stage states, represented by different background colours (brown and violet). One transition was more likely (common; 70% transition probability), the other less so (rare; 30% transition probability). In the second-stage, another two-option choice between stimuli was required, and was reinforced at one of three different outcome levels (referred to as “reward”; “high” is big reward and no delay; “medium” is small reward and small delay; “low” is no reward and big delay; see *SI Methods*). In both decision stages, the choice stimuli were randomized to two of three possible locations. To encourage learning, the outcome level for each second-stage option was dynamic, remaining the same for 5-9 trials, then changed randomly to any of the three possibilities (including remaining the same).

We first assessed MF and MS RL by exploring how the previous trial’s reward and transition type (common or rare) affected current first-stage choice. MF-RL does not exploit information about task structure, so it predicts no difference in the probability of repeating a first-stage choice dependent on the transition (simulations in Fig. S1A). By contrast, the key signature of MS-RL is just such a difference (simulations in Fig. S1B). Both subjects were indeed much more likely to repeat the same first-stage choice if a high reward was obtained through a common transition than when obtained following a rare transition (Fig. 2A). The opposite pattern was seen following either a medium or a low reward.

**Fig. 2.**
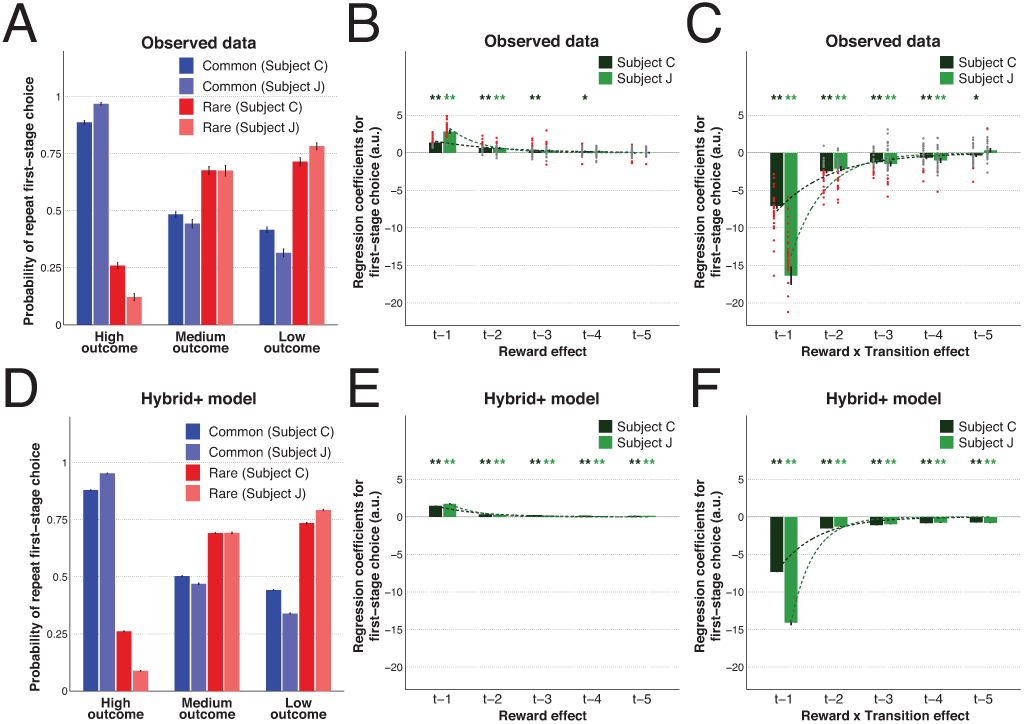
The impact of both reward and transition information on first-stage choice behaviour. (A) Likelihood of first-stage choice repetition, averaged across sessions, as a function of reward and transition on the previous trial. Error bars depict SEM. (B-C) Logistic regression results on first-stage choice with the contributions of the reward main effect (B) and reward × transition (C) from the five previous trials. Dots represent fixed-effects coefficients for each session (red when *p* < 0.05, grey otherwise). (D-F) Similar results obtained from simulations (100 runs per session and respecting the exact reward structure subjects experienced) using the best fit *Hybrid*+ model. Bar and error bar values correspond, respectively, to mixed-effect coefficients and their SE. Dashed lines illustrate the exponential best fit on the mean fixed-effects coefficients of each trial into the past. ** *α* = 0.01 and * *α* = 0.05 in two-tailed one sample t-test with null-hypothesis mean equal to zero for the fixed-effects estimates.

To quantify the influence of MF and MS RL further, we assessed first-stage choices using multiple logistic regression (i.e. aiming to predict the chosen picture at first-stage), taking into account the first-stage choice (C), reward (R) and transition (T) information of up to five trials in the past (Fig. 2B and C; Table S1). For relevant learning rates, a pure MF learner’s choices will chiefly be determined by the reward that the choice on the previous trial delivered (see the R × C predictor in Fig. S2A), whereas those of a pure MS learner will also be influenced by whether the transition was common or rare (see the R × T × C predictor in Fig. S2B). This is because in a MS agent a good reward from a rare transition will enhance the probability of the choice that was not taken, for which that second-stage state is a more likely reward. Choices derived from an agent combining both systems will balance the MF main effect of reward with the MS interaction (Fig. S2C). We found a significant main effect of previous reward on observed first-stage choice repetition (see *t* − 1 effect on Fig. 2B and R_*t*−1_ × C_*t*−1_ on Table S1). In addition, consistent with MS, a significant effect of previous reward × transition was also present, reflecting the adaptive switch in first-stage choice following a high reward obtained through a rare transition (see *t* − 1 effect on Fig. 2C and R_*t*−1_ × T_*t*−1_ × C_*t*−1_ on Table S1). Moreover, these two predictors were not only both significantly different from zero (both subjects fixed-effects *F*-tests *p* < 0.001 in all sessions; mixed-effects *F*(2) = 386.07/173.68, *p* < 0.001/< 0.001 for C/J), but the weight of the reward × transition interaction was significantly greater than the main effect of reward (both subjects fixed-effects *F*-tests, with *p* < 0.001 in all sessions; mixed-effects *F*(2) = 577.68/231.14, *p* < 0.001/< 0.001 for C/J), indicative of greater reliance on MS-RL. Finally, as has previously been noted (8, 23), both subjects tended to perseverate on the same first-stage choice irrespective of any other variable (*p* < 0.001 for both subjects; see predictor C_*t*−1_ on Table S1).

According to both MS and MF RL, the effects of trials in the further past tail off, typically exponentially (1). We found that the contribution to first-stage choice from both reward history (Fig. 2B) and combined reward × transition information (Fig. 2C) reduced across five trials into the past in a way consistent with an exponential decay fit (decay constants of reward −0.78/-1.62, adjusted R^2^ = 0.46/0.69; decay constants of reward × transition −0.94/-1.50, adjusted R^2^ = 0.71/0.82 for C/J). Despite this decay, these MS and MF RL effects on current choice were present in each of the five trials into the past (Table S1). Overall, our logistic analysis indicated both MF and MS RL strategies coexist, but MS-RL had significantly greater influence over choice in both subjects.

### Computational modelling results

To validate and enrich the logistic regression analysis, we fitted a variety of pure MF (Tables S2 and S3; Fig. S1, S2 and S3) and pure MS RL models (Table S4; Fig. S1, S2 and S3) to each subject’s trial-by-trial choices using both fixed-effects (individual fits for each session) and mixed-effects (taking parameters of each subject as random effects across sessions) fitting procedures. As in previous studies (8, 24), we also considered a *Hybrid* model in which the best MF and MS models operated in parallel, and with their decision values being combined to determine choice probabilities (Table S5; Fig. S1, S2 and S3). This uses a parameter (*ω* ∈ [0, 1]) for the relative weight of MS (*ω* ≃ 1) and MF (*ω* ≃ 0) control (8, 25). A careful examination of the data revealed that this *Hybrid* model required further refinement, leading us to develop a novel *Hybrid*+ model (see below), which accurately reproduced the strong influence of the previous trial reward on current choice.

The complexity-adjusted likelihoods of the models were compared to determine which best fit the behavioural data (Table S6). In both subjects, choice behaviour was best explained by a combined MF and MS strategy, corroborating our logistic analysis. The best *Hybrid* model fit had a lower *BIC* score and a higher exceedance probability than the best pure MF and best pure MS models. This winning approach combined the *SARSA* MF model (a better pure MF approach than *Q*-learning) without an eligibility trace parameter, and the *Forward*_1_ MS model (the best pure MS approach) for which the state-transition probabilities are assumed known from the beginning of the task (see *SI Methods* for explanation of differences between each of the MF and MS models).

With regard to the balance between MF and MS control (Table 1), the mean of the *ω* hyperparameter was close to 90% in both subjects (different from 0 and 100% with *p* < 0.001 on sign tests in all sessions), in line with the MS dominance found in our regression analysis. The best-fit learning rate, *α*, was the same for both decision stages and was relatively high (close to 0.8 in both subjects, Table 1), probably due to the non-stationary and occasionally switching second-stage reward structure. On the other hand, first-stage choice was more deterministic than second-stage choice (*β*_1_ > *β*_2_, Table 1). Finally, the modeling also captured the small but positive tendency to repeat recently chosen options (parameter *κ*, Table 1).

**Table 1.**
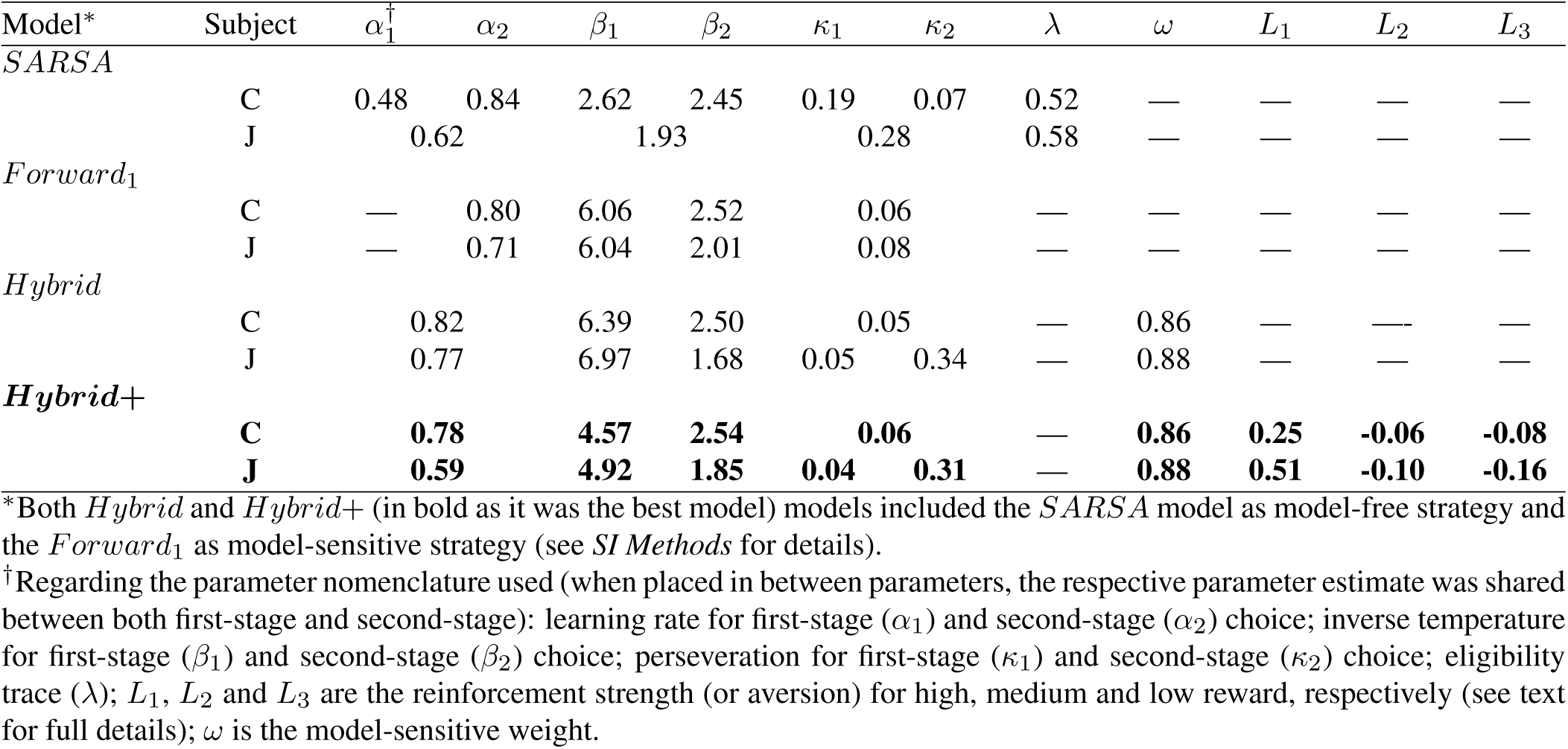
Best fitting mixed-effects hyperparameters from the best models of each reinforcement learning approach.

### Model validation and simulation results

An important test for the models concerns whether they can accurately replicate the observed choice behaviour. Therefore, we used the best RL models for each learning strategy to simulate choice data on the same task, and then analysed the resulting simulated behaviour in the same way (Figs. 2D-F, S1, S2 and S3). These generated data confirmed the previously described differences between MF and MS RL and confirmed the qualitative validity of the best *Hybrid* model. However, they also highlighted important quantitative limitations. One of the most striking differences between the *Hybrid* model simulations and the observed data was the excess weight given to the most recent trial and, consequently, the discrepancies in the exponential decays (compare Fig. 2B-C with Fig. S3C; decay constants for the reward main effect observed −0.78/-1.62 versus simulated −0.37/-0.36 for C/J; and reward × transition effect observed −0.94/-1.50 versus simulated −0.22/-0.17 for C/J).

This overweighting of the previous trial is akin to a sophisticated, MS, form of perseveration – i.e., a one step credit assignment influence on choice depending on reward and transition information of the last trial. For pure perseveration, the *Q*_*Hybrid*_ value of the previous first-stage choice is boosted, independent of the transition or reward. For this new effect, the influence on the *Q*_*Hybrid*_ value of the previous first-stage choice could depend on both reward and transition, with a factor dependent on the outcome level of the previous trial (*L*_1_, *L*_2_ or *L*_3_, for high, medium and low) being added or subtracted according to whether the transition on the previous trial was common or rare, respectively. This way, a positive value (*L* > 0) denotes the strength of the reinforcement by reward, whereas a negative value (*L* < 0) quantifies the aversion for that particular reward. We call this new model *Hybrid*+.

Comparisons (Table S6) showed that *Hybrid*+ outperformed all the other RL accounts, including pure MS reasoning and the previous *Hybrid* model (all exceedance probability values > 0.99). The extra parameters were justified according to the *BIC, BIC*_*int*_ and the exceedance probability. An important question is whether the original RL parameters remained stable after refitting all parameters. Indeed, very few changes were observed (Table 1). Critically, *Hybrid*+ captured behavioural characteristics that eluded *Hybrid*; the simulated choice data generated by *Hybrid*+ successfully captured not only the observed pattern of repeat probability at first-stage choice (Fig. 2A and 2D), but also the profiles of both reward main effect (Fig. 2B and 2E) and reward × transition interaction (Fig. 2C and 2F) shown in the logistic regressions. Moreover, the best-fitted values of the additional parameters (*L*_1_, *L*_2_, *L*_3_) revealed that high reward had a high reinforcement strength, but both medium and low reward had an aversive impact (Table 1), as previously noted. Thus, both model comparison and simulation results supported the validity of the new *Hybrid*+ account.

To examine the relationship between both descriptive and computational results, we explicitly compared coefficients obtained from the regression with the best fit *Hybrid*+ parameters for each subject and session. In addition, we also simulated new data from *Hybrid*+ using those parameters, performed logistic regression on these new data, and compared the resulting coefficients with the generating parameters. We found that stronger (i.e., more negative) reward × transition interaction effects were associated with greater MS *ω Hybrid*+ parameters (Fig. S5A). We also observed a significant negative correlation between the first-stage inverse temperature parameters (lower values reflect stochasticity in choice) and the residuals from the regression (Fig. S5B), and a positive correlation between both logistic and computational first-stage choice perseverance measures (Fig. S5C). Taken together, these results demonstrate the strong correspondence between the regression analysis and computational modelling approaches.

Finally, to test the tradeoff and stability of MF vs. MS influences over time, as well as whether habits were forming with repeated experience of the task (4), we assessed the correlations between model parameters and their respective session number (Fig. S6). We found no significant relationship between session number and the main effect of previous reward, nor a reduction in the *ω* parameter across sessions (Fig. S6A-B). On the other hand, the previous reward × transition interaction effect reduced across sessions in both subjects (Fig. S6C).

This effect is likely caused by enhanced stochasticity, associated with a reduction in the first-stage inverse temperature parameter (Fig. S7), which would be expected to reduce the logistic regression weights. Another is the one-step MS contributions in the additional parameters of *Hybrid*+: we found the *L*_1_ parameter decreased across sessions (Fig. S8), implying the influence of the high reward is reduced. Taken together, these results highlight the potential advantage of the computational modelling approach (over the regression approach) to decouple structurally different contributors (i.e., perseveration and MF/MS weight) that may otherwise be captured in a single regression weight (reward × transition).

### Reaction time analysis

Various (and potentially conflicting) considerations might affect reaction times (RTs), including the expectation that changing a choice might take longer than repeating one, and the observation that first-stage choices can be planned as soon as the reward is revealed on the previous trial, whereas second-stage choices cannot, since they depend on the transition (albeit not in the conditions employed by (26)). We therefore analyzed RTs from a number of perspectives.

First-stage RTs were fast, and significantly shorter than second-stage RTs (first-stage: mean±sd = 499±201 for subject C and 647±191ms for subject J; second-stage mean±sd = 514±210 for subject C and 663±194ms for subject J; *p* < 0.001 for both two-sample t-tests). In both subjects, we found consistent RT differences between common and rare trials as a function of high or low rewards received (Fig. 3). First-stage RTs were slower following high rewards obtained through rare versus common transitions, and low rewards obtained through common versus rare transitions. When considered alongside choice data (Fig. 2A), these slower RTs occurred when the likelihood of choice switching was highest, and were just the situations when model sensitivity is most acute.

**Fig. 3.**
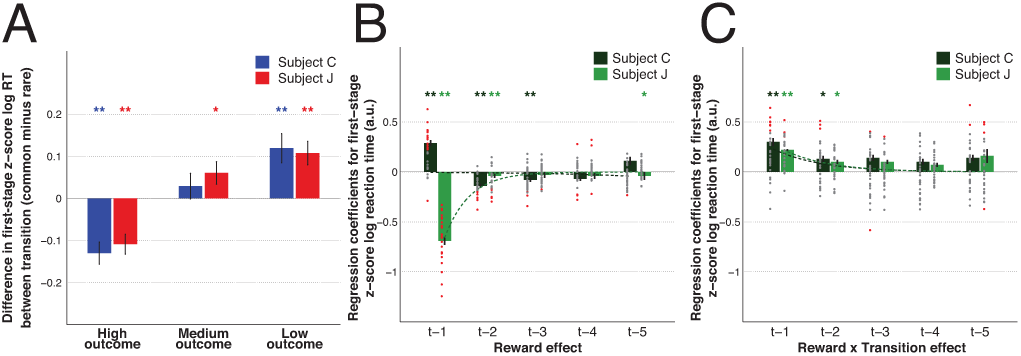
The impact of both reward and transition information on first-stage choice reaction time. (A) The averaged across sessions *z*-scored first-stage reaction time (RT) difference between previous common and previous rare trials as a function of reward on the previous trial (high *z*-scores indicate responses faster if previous transition was rare). Error bars depict SEM. (B-C) Multiple linear regression results on first-stage reaction time with the contributions of the reward main effect (B) and the reward × transition interaction term (C) from the five previous trials. Dots represent the fixed-effects coefficients for each session (coloured red when *p* < 0.05 and grey otherwise). Bar and error bar values correspond, respectively, to the mixed-effect coefficients and their SE. Dashed lines illustrate the exponential best fit on the mean fixed-effects coefficients of each trial into the past. ** *α* = 0.01 and * *α* = 0.05 in two-tailed one sample t-test with null-hypothesis mean equal to zero.

We also performed multiple linear regression on first-stage RT (Fig. 3 and Table S7). Despite some similarities with the approach used for choice behaviour, we note that in our predictive model for first-stage RT the effects of previous reward, transition and reward × transition do not include the interaction with previous first-stage choice information. In both subjects, the RT was modulated over multiple past trials by both the reward (Fig. 3B) and the reward × transition (Fig. 3C). Although the interaction term was similar between subjects, the effect of previous reward differed: a high reward (independent of transition) on trial *t* − 1 led to faster/slower RTs in subject J/C, respectively (also seen in Fig. 3A). These *t* − 1 RT differences might be explained by differential trial lengths between subjects (delays and task timings were shorter for C than J to maintain motivation) or by different speed/accuracy strategies following high rewards.

A different index of response vigour and task motivation is how quickly subjects acquire fixation (fixRT) based on previous reward. Both subjects exhibited a main effect of reward on fixRT (i.e., faster fixation following better rewards; Fig. S9A). No effect of the reward × transition was found (Fig. S9B), suggesting that fixRT reflected only a MF influence, whereas RT was influenced by both MF and MS systems.

## Discussion

There is now a large body of work on the combination of MF and MS influences, and a collection of forms of MS reasoning, even just in the various versions of the task we studied (8, 10, 14, 16, 27–29). In addition to revealing fundamental features of behavioural strategies, changes in MS influences have been associated with various psychiatric (30–33), neurological (34) and genetic, pharmacological or stimulation-induced manipulations (35–38). It is therefore pressing to examine more closely the various components of behavior. Our behavioural and computational results demonstrate that, like humans performing an equivalent RL task (8), non-human primates employ both MF and MS RL strategies. In our subjects, reward history (relevant for both learning strategies) and state-transition knowledge (used in MS computations) had a significant impact on choice, and such influence decayed exponentially as a function of trials into the past. This was evident both via logistic regression and RL-based analyses. MS-RL comparatively dominated, and we demonstrated that such MS dominance is stable over extended experience of a task.

We validated our analyses by extracting the same summary statistics and performing the same regression analyses on choices generated from the best-fitting models as on the actual data (39). This helped elucidate the role played by the RL parameters, being evident, for instance, in the correlation between the reward × transition interaction coefficient and the MS weight parameter (*ω*). More importantly, this highlighted residual structure in the actual data that was not evident in the data generated from the original *Hybrid* model (8), suggesting further model refinement was necessary.

In particular, an excessive influence of the immediately previous trial motivated the novel *Hybrid*+ model, which closely reproduced the observed choice behaviour. Its extra parameters changed the influence of the action chosen on that trial as a function of the reward and the transition. We considered this to be a form of sophisticated, MS perseveration, effectively interpreting it as a MS influence over MF evaluation coming from a one-step working-memory-for-state representation (15). Such an influence is MF, since it depends on a direct effect of the past trial rather than an assessment of a future one; however, the way it differentiates common and rare transitions makes it MS. Other limited MS strategies have been noted in the case of serial reversals (40). MS perseveration could also be seen as a short term form of sophisticated counterfactual or regret/rejoice-based influence on the unchosen action (41), and so be added to other MF modulations such as forgetting (42). Most critically, model comparison showed that this effect co-existed with conventional MS and MF reasoning – implying that it resolved a significant problem with the fit of the data.

Within the hybrid models, we examined different possibilities for the MF component. We found that *SARSA* (which evaluates the second-stage according to the estimated value of the choice the subject actually took on a trial) fit better than *Q*-learning (which uses the value of the better of the two choices). This is consistent with previous reports in non-human primates performing a very different task (18), though evidence from rodents favours *Q*-learning (19). It was also notable that the best-fitting MF model involved no eligibility trace (i.e., *λ* = 0); this implies, for instance, that it takes the MF component at least two trials to change its estimate of the value of a first-stage action following a change after the second-stage.

In keeping with the observation that many manipulations reduce MS control without increasing MF control, it has been suggested that MF influences might instead arise from MS reasoning with incorrect or incompetent models (43). The MS perseveration effect could perhaps be seen as an example of this. There is, of course, a large range of possible flawed models; however, the extensive training (and the large value of *ω* we found) perhaps suggest that this problem might be less severe in our study.

The best-fitting MS-RL strategy (excluding the MS perseveration effect) treated the state-transition probabilities as being known from the start, which is consistent with the extensive training the subjects ultimately received. That MS control dominated more here (*ω* near 90%) than in recent human studies (*ω* approximately 40-60%; (8, 36); or *ω* → 0; (24)) could result from the non-stationarity of the outcomes (changing every 5-9 trials), which should optimally favor the more flexible, MS, controller (6) or be another effect of the extensive training, as also seen in human studies (16). It could arise from an increase in the efficiency of the implementation of MS reasoning, for instance from reducing its computational cost and increasing its speed. Either of these might come by arranging for a progressively greater MF implementation of MB reasoning via representational change (15), as we argued above for the MS perseveration effect.

Theoretical accounts have suggested a speed accuracy trade-off between MF and MS computations, with the former being fast and at least explicit versions of the latter relatively slow (44, 45). Indeed, first-stage RT analysis confirmed that decisions that showed sensitivity to both reward and transition structure took longer. This RT effect followed a similar exponential decay with trials into the past as in the choice data. It would be harder to square with the suggestion that faster responses arise from the chunking of sequential actions (26), something that our design deters, with randomized positions for second-stage stimuli (8). It also militates against MS proposals emphasizing pre-computations at the time of outcome, where the re-evaluation of the utility of states given the received rewards helps future choice (10, 21, 46). Overall, the RT evidence is supportive of a forward looking MS valuation process happening at the time of choice (47, 48), as in the original conception of MB reasoning in this task (8).

It was notable that the RTs, particularly the fixRT, were more strongly influenced by the main effect of reward, than any effect of transition or reward × transition. This may be consistent with the observation that the average reward rate, estimated in a MF way from recent past trials, and putatively reported through tonic activity of dopamine neurons, is a main mediator of the vigor of actions (49–51).

In conclusion, we have been able to show clear evidence of combined MF and MS RL behaviour in non-human primates. Our computational analyses of choice suggested an enriched picture of the combination; the analyses of RTs showed that they are subject to different influences. Future studies focusing on the neural signals may uncover the biological substrates of these computational mechanisms.

## ACKNOWLEDGEMENTS

B.M. was supported by the Fundacão para a Ciência e Tecnologia (scholarship SFRH/BD/51711/2011). N.M. was supported by Astor Foundation, Rostrees Charitable Trust. T.E.J.B. was supported by a Wellcome Trust Senior Research Fellowship (WT104765MA) and funding from the James S McDonnell Foundation (JSMF220020372). P.D. was supported by The Gatsby Charitable Foundation. S.W.K. was supported by a Wellcome Trust New Investigator Award (096689/Z/11/Z). The authors thank Thomas Akam, James Butler and Tim Muller for useful discussions. PD is now at the Max Planck Institute for Biological Cybernetics, Tübingen.

## SI Methods

### Subjects and experimental apparatus

Two rhesus monkeys *Macaca mullata* were used as subjects: subject C weighing 8 Kg; and subject J weighing 11 Kg. Daily fluid intake was regulated to maintain motivation on the task. During the experiment, subjects were seated in a primate chair inside a darkened room with their heads fixed and facing a 19-inch computer screen (60Hz video refresh rate) positioned 62 cm from the subject’s eyes. Each subject’s eye position and pupil dilation was monitored with an infrared eye tracking system having a sampling rate of 240 Hz (ISCAN ETL-200). Both subjects indicated their choice by moving a joystick with a left arm movement towards one of three possible locations (C: left, right and down; J: left, right and up). The reward (C: cranberry juice diluted to one-fourth with water; J: apple juice diluted to one half with water) was provided by a spout positioned in front of the subject’s mouth and delivered at a constant flow-rate using a peristaltic pump (Ismatec IPC). We used Monkeylogic software (http://www.monkeylogic.net/): to control the presentation of stimuli and task contingencies; to generate timestamps of behaviourally-relevant events; and to acquire joystick as well as eye data (1000 Hz of analog data acquisition). All visual stimuli used were the same across sessions for both subjects, and were presented at pre-determined degrees of visual angle (see below). Six decision option pictures were chosen from a stimulus database, reduced in size and modified through a custom-made image processing algorithm to make the average luminance equivalent for all. Similarly, the background colours used (grey, violet and brown) were tested with a luminance meter and adjusted accordingly. Finally, three stimuli used as secondary reinforcers were generated as different spatial combinations of the same number of dark pixels in a white background, also to assure luminance equality. All experimental procedures were approved by the UCL Local Ethical Procedures Committee and the UK Home Office, and carried out in accordance with the UK Animals (Scientific Procedures) Act.

### Task: design and timeline

Subjects performed a two-stage Markov decision task (see Fig. 1), similar to the one used in a previous human study (8) that was designed to detect simultaneous signatures of MF and MS systems as they concurrently learn. In brief, two decisions had to be made before the subject received an outcome (see Fig. 1A). The first-stage state was represented by a grey background and the choice was between two options presented as pictures (the same fixed set of pictures was used throughout the entire task). Each of these first-stage choices could lead to either a common (70% transition probability) or rare (30% transition probability) second-stage state, represented by different background colours (brown and violet). This state-transition structure was kept fixed throughout the experiment. In the second-stage, another two-option choice between stimuli was required and it was reinforced according to one of three different levels of outcome (see Fig. 1B). Importantly, to encourage learning, each of the four second stage options had independent reward structures according to a form of random walk that was sampled afresh on each session (see below). In both decision stages, each choice option (or each presented stimuli) could randomly assume one of three possible locations (C: left, right and down; J: left, right and up). No significant preference for any first-stage stimulus across sessions was found (both one-sample t-tests with *p* > 0.05) but, given the three physically possible actions, small side biases were observed (both one-way ANOVAs with *p* < 0.01). Fifteen percent of the trials were forced, i.e., where only one stimulus was presented – these could be at either the first or second-stage. Unless stated otherwise, such forced trials were not included in the data analysis. The trial type sequence was randomly generated at the start of the session and was followed even after error trials. Error types included trials with no choice, no eye fixation, eye fixation break, early joystick response, joystick not centred before choice or movement towards a location not available. Error trials resulted in time-outs for the subjects. Unless otherwise specified, we excluded such trials from the data analysis (C: *M* = 5%; J: *M* = 8%).

The outcome (referred to as “Reward”) could assume one of three categorical levels, defined according to the amount of juice delivered (determined by the time the juice pump was on) and a specific delay (in addition to a fixed 500 ms delay common to all outcome levels) before juice delivery. Therefore, the reward could be: high (big reward and no delay), medium (small reward and small delay) or low (no reward and big delay). The precise reward amounts for big and small rewards were tailored for each subject to ensure that they received their daily fluid allotment over the course of the experimental sessions. Consequently, the duration for which the reward pump was active (and hence the magnitude of delivered rewards) differed slightly between the two subjects. Furthermore, instead of a fixed reward amount, big and small rewards corresponded to non-overlapping time intervals (C: high reward ranged on average from 682 to 962 ms and medium reward ranged on average from 117 to 390 ms; J: high reward ranged on average from 976 to 1257 ms and medium reward level ranged on average from 507 to 826 ms) of juice delivery where a small Gaussian drift (mean/standard deviation of 0/200 ms for high reward and 0/100 ms for medium reward) was added. This was used not only to promote constant valuation of the reward amount, but also to help the computational model fitting procedure. The additional specific delay periods were fixed throughout the experiment but varied across subjects (C: 750 ms for small delay and 2500 ms for big delay; J: 1500 ms for small delay and 4000 ms for big delay). Importantly, for each of the second-stage pictures the outcome level remained the same for a minimum number of trials (a uniformly distributed pseudorandom integer between 5 and 9) and then, either stayed in the same level (with one-third probability) or changed randomly to one of the other two possible outcome levels. Three different stimuli were used as secondary reinforcers, providing feedback for each of the three outcome levels. Both subjects had prior classical conditioning training with these stimuli (see Fig. 1B), with the above mentioned reward magnitude ranges and delays for each outcome level used in the experiment being respected.

The sequence of events in the behavioural task is shown in Fig. 1A. Each trial started with the presentation of a grey background (start epoch). A central square fixation cue 0.4°in width then appeared after a random interval of 200-500 ms. After this, subjects were required to keep the joystick in the centre position as well as to maintain eye fixation within 3.4°(C) or 2.8°(J) of the cue for a 500 ms (C) or 750 ms (J) period (fixation epoch). Then, the fixation cue was removed and two stimuli (5°in size) appeared at 7°away from fixation in the available locations (choice epoch). During the task, in the absence of a fixation cue, the animal was free to look around. The maximum time allowed for eye fixation as well as response with the joystick was 5000 ms for both choice stages. After a choice was made, the non-selected stimulus was removed and the background color changed according to the second-stage state to which the transition had occurred (transition epoch). After 500 ms, the stimulus selected in the first-stage was removed from the screen. Similar fixation and choice epochs were used for the second-stage. Once the choice had been made in the second-stage, the non selected stimulus was removed and the selected one remained for 750 ms before the secondary reinforcer stimulus (5°square) appeared at the center of the screen (pre-feedback epoch). Following its appearance, the feedback stimulus remained present for 750 ms. After the removal from the screen of the secondary reinforcer, a fixed 500 ms delay period occurred before either the reward delivery (for high reward) or both small and big additional delays started (for both medium and low rewards, respectively). Therefore, a total of 1250 ms was the minimum time from the secondary reinforcer presentation to the delivery of any juice (feedback epoch). The inter-trial period duration was 1500 ms (ITI epoch).

### Behavioural analysis

All analyses were conducted using MATLAB^©^ R2014b (MathWorks). Statistical significance was assessed at *α*=0.05, unless otherwise stated. Behavioural variables were defined as: C is first-stage choice (1=car picture, 0=watering can picture); R is outcome level (referred to as “Reward”; assumed as continuous, with low=1, medium=2, high=3); and T is transition (rare=1, common=0). In regressions, these variables were mean centred, and continuous variables were also scaled by dividing them by twice their standard deviations so that the magnitudes of regression coefficients could be directly compared (52). To quantify the factors predicting first-stage choice at trial *t*, C_*t*_ a multiple logistic regression was used in which the predictors included information from the last 5 trials, *i* ∈ {1,2,3,4,5}, and were: Const (constant term) captured any potential first-stage picture bias; C_*t*−*i*_, modelling a potential independent tendency to stick with the same option; R_*t*−*i*_, T_*t*−*i*_, R_*t*−*i*_ × T_*t*−*i*_, measuring any potential preference in first-stage picture choice given the previous reward, the previous transitions and the interaction effect of both, respectively; R_*t*−*i*_× C_*t*−*i*_, T_*t*−*i*_× C_*t*−*i*_, R_*t*−*i*_×T_*t*−*i*_× C_*t*−*i*_, were the predictors of interest which quantified the main effects of reward, transition and the reward × transition interaction effect, respectively. Although unexpected, both subjects showed a small but significant main effect of transition (Fig. S4A) but a similar effect was present in the simulations derived from our best RL model (Fig. S4B) suggesting that correlations within the task design and reward structure may underlie this effect. Linear hypothesis testing on the vector of regression coefficients (performed for each individual session in the fixed-effects; and using the estimated mixed effects for each predictor) was performed to test either if more than one coefficient or a difference between coefficients was significantly different from zero. First-stage RT was defined as the time from first-stage stimuli presentation to joystick movement towards the specified location (all side locations with the same target radius). For each subject and session, first-stage RT were independently *log* transformed and *z*-scored for the three possible side responses (this was done as side RT differences were with both one-way ANOVAs with *p* < 0.001). Data points greater than three times the SDs from the individual means were removed. The first-stage eye fixation time (fixRT) was defined as the time from fixation cue presentation to the first time the x and y position eye position waswithin that subject’s required fixation radius. The raw data was then *log* transformed and *z*-scored. To determine the effect of behavioural variables on first-stage RT and fixRT, we performed a multiple linear regression analysis on the current trial *t log* transformed and *z*-scored first-stage RT/fixRT, using as predictors: F_*t*_, used to model (linearly-increasing) fatigue by counting the trials in the session; R_*t*−*i*_, T_*t*−*i*_ and R_*t*−*i*_× T_*t*−*i*_ were the predictors of interest which quantified the main effect of reward, the main effect of transition and the reward × transition interaction effect, respectively.

### Regression analysis fitting

Fixed-effects (fitting the regression models individually to each session) and mixed-effects (assuming regression coefficients to be random effects across sessions) analyses were performed for each subject. Fixed-effects fitting was performed using a generalized linear model regression package (`glmfit` in MATLAB with: a binomial distribution and the logit link function for logistic regressions, a normal distribution and the identity link function for linear regressions), and the statistical importance of each predictor’s estimates was assessed by both the p-values obtained from each session as well as their distribution across sessions (two-tailed one-sample t-test for a mean of 0 and unknown variance). Mixed-effects fitting was achieved with either a non-linear model with a stochastic approximation expectation-maximization method for logistic regression (`nlmefitsa` in MATLAB with importance sampling for approximating the loglikelihood) or a linear model method for the RTs (`filme` in MATLAB). The standard errors for the coefficient estimates as well as their 95% confidence intervals (CI) were reported.

### Computational modelling

We fitted choice behaviour in the task in a similar manner to previous human studies (53), assessing three different reinforcement learning approaches: MF learning, MS learning and a hybrid strategy combining the decision values of both (8, 24). The task consists of three states (first stage: *A*; second stage: *B* and *C*), each with two actions (*x* and *y*). Importantly, we assume that the subjects already know that the action corresponds to the choice of a picture belonging to the respective state (rather than the side, given their very modest side biases). The main goal is to learn to compute a state-action value function, *Q*(*s, a*), mapping each state-action pair to its expected future value. On trial *t*, the first-stage state (always *s*_*A*_) is denoted by *s*_1,*t*_, the second-stage state by *s*_2,*t*_, the first and second-stage actions by *a*_1,*t*_ and *a*_2,*t*_ and the first and second-stage rewards as *r*_1,*t*_ (always zero) and *r*_2,*t*_. For the model fitting *r*_2,*t*_ corresponded to the amount of juice delivered at trial *t* divided by the maximum amount of juice obtained by the subject within the entire respective session.

In MF-RL the value for the visited state-action pair at each stage *i* and trial *t, Q*(*s*_*i,t*_, *a*_*i,t*_), is updated based on the temporal difference prediction error, *δ*_*i,t*_, which sums the actual reward *r*_*i,t*_ and the difference between predictions at successive states *s*_*i*+1,*t*_ and *s*_*i,t*_. For the first-stage choice, *r*_1,*t*_ = 0 and *δ*_1,*t*_ is driven by the second-stage value *Q*(*s*_2,*t*_, *a*_2,*t*_). On the other hand, at second-stage there is no further value apart from the immediate reward, *r*_2,*t*_, and ultimately the start of a new trial. For convenience, we create a fictitious state, *s*_3,*t*_, and action, *a*_3,*t*_, for which *Q*(*s*_3,*t*_, *a*_3,*t*_) is always 0. Two different MF-RL models were used to fit behaviour: the *SARSA* variant of temporal difference learning (54), which has previously been observed in non-human primates (18); and the *Q*-learning model, as described in rodents (19).

In *SARSA*, state *s*_*i*+1,*t*_ is evaluated according to the actual action *a*_*i*+1,*t*_ that the subject selects. This makes the prediction error:

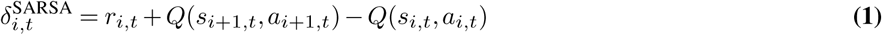

By contrast, in *Q*-learning, the state is evaluated based on what the subject believes to be the best action available there, independent of the policy being followed. This makes the prediction error:

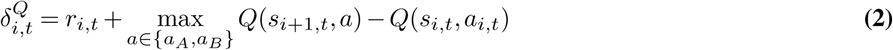

Either of these errors in the estimate drives learning by correcting the respective MF prediction through the following update rule:

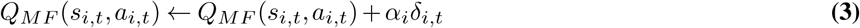

where *α*_*i*_ is the learning rate at stage *i*, and was fit to the observed behaviour. In previous work, different learning rates were found for the stages (8). Given the two-stage design of the task, the model also permits an additional stage-skipping update of first-stage values by having an eligibility trace parameter *λ* (1), which connects the two stages and allows the reward prediction error at the second-stage to influence first-stage values:

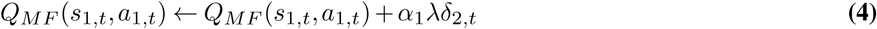

The parameter *λ* was also fit to the observed behaviour. Consistent with the episodic structure of the task (with an explicit inter-trial epoch), it is assumed that eligibility does not carry over from trial to trial.

In MS-RL, the agent not only maps state-action pairs to a probability distribution over the subsequent state but also learns the immediate reward values for each state. More specifically, it requires knowledge of the probabilities with which each first-stage action leads to each second-stage state, as well as learning the expected reward associated with each second-stage actions. The MS second-stage state-action values *Q*_*MS*_ (*s*_2,*t*_, *a*_2,*t*_) are just estimates of the immediate reward *r*_2,*t*_, and so coincide with MF values there (since *Q*(*s*_3,*t*_, *a*_3,*t*_) = 0). We define *Q*_*MS*_ = *Q*_*MF*_ at those states. On the other hand, the first-stage action values *Q*_*MS*_ (*s*_1,*t*_, *a*_1,*t*_) differ and are computed by weighting the estimates on trial *t* of the rewards by the appropriate probabilities:

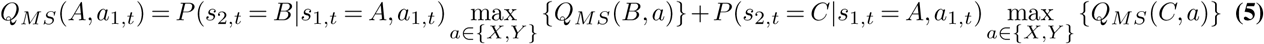

Different approaches to estimating the state-transition probabilities give rise to three different MS models, designated here as *Forward*_1_, *Forward*_2_ and *Forward*_3_. In the first model, the agent had explicit knowledge of the correct state-transition probabilities, *P* = {0.3,0.7}. The extensive training of both subjects prior to this experiment makes this plausible. In the second model, agents were assumed to map action-state pairs *a*_1_, *s*_2_ to transition probabilities, *P* = {0.3,0.7}, by counting whether they had more often encountered transitions *a*_1_ = 1, *s*_*B*_ and *a*_1_ = 2, *s*_*B*_ or transitions *a*_1_ = 1, *s*_*C*_ and *a*_1_ = 2, *s*_*B*_ and concluding that the more frequent category corresponds to *p* = 0.7. This latter model corresponds to the one used in the modelling of the original two-step task study (8). Finally, in the *Forward*_3_ model the agent incrementally learn the transition structure by performing an hypothesis test between *p* = { 0.3,0.7} versus *p* = {0.5,0.5} with an additional parameter (*ζ*) modelling the weight given to each of these models. In both *Forward*_2_ and *Forward*_3_ the data for the hypothesis test was reset at the start of every session.

Finally, a so-called *Hybrid* model assumes that first-stage choices are computed as a weighted sum of the state-action values from MF and MS learning systems:

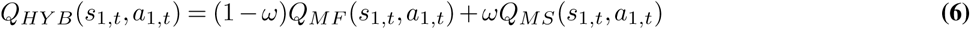

where *ω* is a weighting parameter that determines the relative contribution of MS and MF values. When *ω* = 0 the model reflects pure MF control; when *ω* = 1, it reflects pure MS control. For convenience the hybrid model was constructed using the best fitting MF (*SARSA* model) and MS (*Forward*_1_) models, given the computational burden of fitting all possible combinations simultaneously.

A careful examination of the data revealed that the original hybrid model required further refinement in order to reproduce more accurately the strong influence of the previous trial on the present one. In this new *Hybrid*+ model, the value of the chosen (*a*_1,*t*_) or unchosen (*a* ≠ *a*_1,*t*_) first-stage action was boosted or suppressed as a function of whether the state-transition (*Trans*) observed at trial *t* was common or rare and the level of the outcome achieved (*Rew*). Algorithmically, after the previously described *Q*_*HY B*_ calculation (Eq. 6) an additional boost (or decrease) occurred according to:

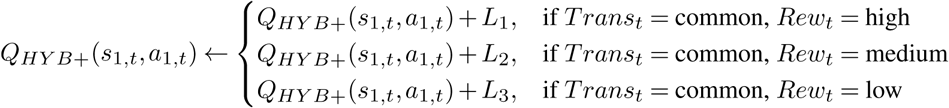

and

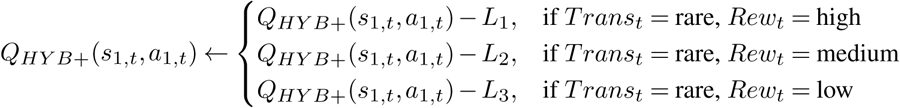

where there are separate parameters *L*_*j*_ for each outcome level which can be positive or negative, expressing support or opposition for that particular outcome level. This extra factor can be seen as a MF implementation of a MS effect (15) – MF, since it depends on an effect of the past trial rather than an assessment of a future one; MS, since it includes a one-step version of the interaction to which MS reasoning leads.

For any of the above reinforcement learning strategies, actions were assumed to be stochastic and chosen for each stage according to action probabilities determined by the respective *Q*-action values:

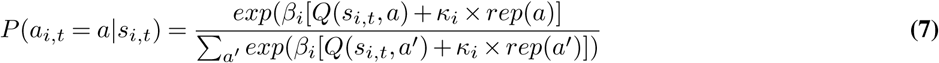

where *β*_*i*_ is the inverse temperature parameter (distinct inverse temperatures are considered for each stage) controlling the determinism of the choices, and so capturing noise and exploration (for *β*_*i*_ = 0 choices are fully random and for *β*_*i*_ = ∞, choices are fully deterministic in the sense that higher-valued options are always preferred). *rep*(*a*) is an indicator variable coding whether the current choice is the same as the one chosen on the previous visit to the same state, with *κ*_*i*_ being a further parameter that captures choice perseveration (*κ*_*i*_ > 0) or switching (*κ*_*i*_ < 0) (23), again with the possibility of distinct values for first and second-stage choices.

In the most general form, the conventional *Hybrid* model involved a total of eight free parameters (*θ* = {*α*_1_, *α*_2_, *β*_1_, *β*_2_, *κ*_1_, *κ*_2_, *λ, ω*}), nesting pure MS (*ω* = 1, with arbitrary *α*_1_ and *λ*) and MF (*ω* = 0) learning as special cases. The *Hybrid*+ model involved three additional parameters *L*_1_, *L*_2_, *L*_3_. We also generated several simpler variants of these models by allowing *α*_1_ = *α*_2_, *β*_1_ = *β*_2_, *κ*_1_ = *κ*_2_, *κ*_1_ = 0, *κ*_2_ = 0 and *λ* = 0. All parameters were fixed within a session, but could vary across sessions.

---

**Fig. S1.**
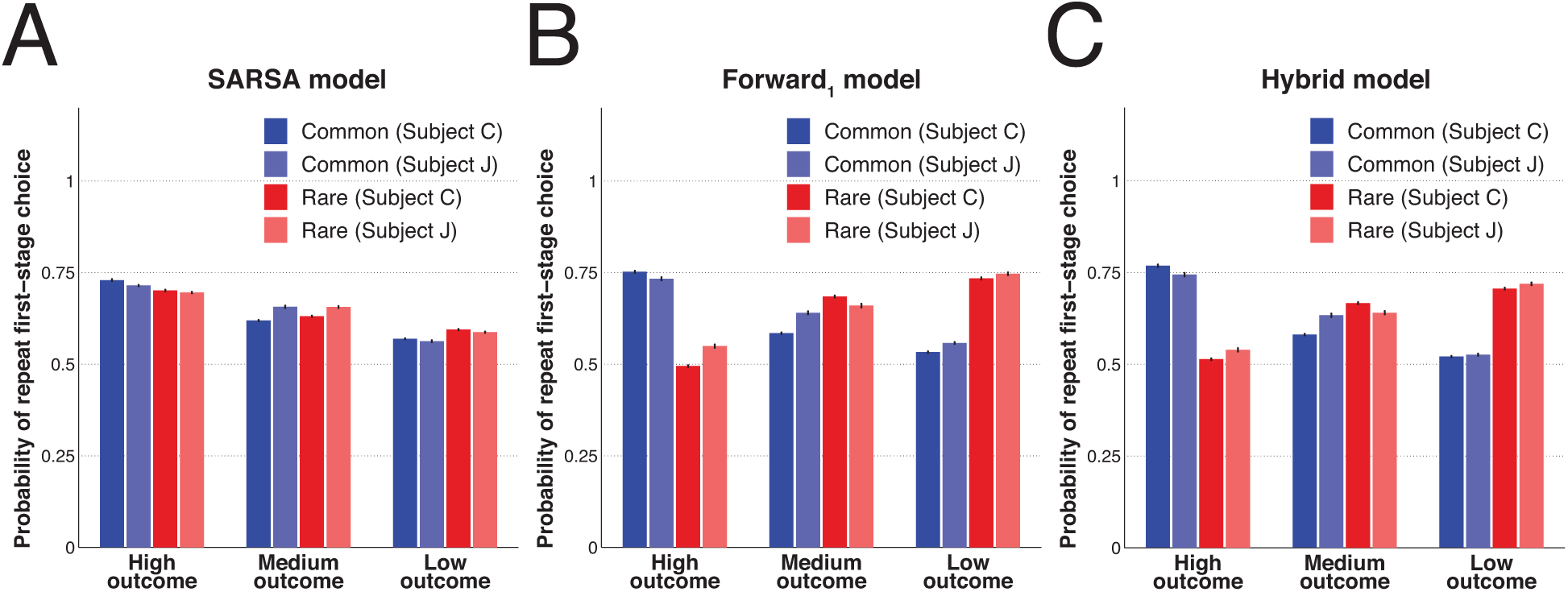
Comparison of the impact of both reward and transition information on first-stage simulated behaviour from each learning strategy. Simulated repetition probabilities as a function of outcome level and transition type for the best pure model-free *SARSA* model (A), the best pure model-sensitive *Forward*_1_ model (B) and the best *Hybrid* model (C). Values were averaged across all sessions, and across 100 simulation runs for each session using the parameters best fit to each subject’s data within each class of model (and respecting the exact same reward structure). Error bars depict SEM.

**Fig. S2.**
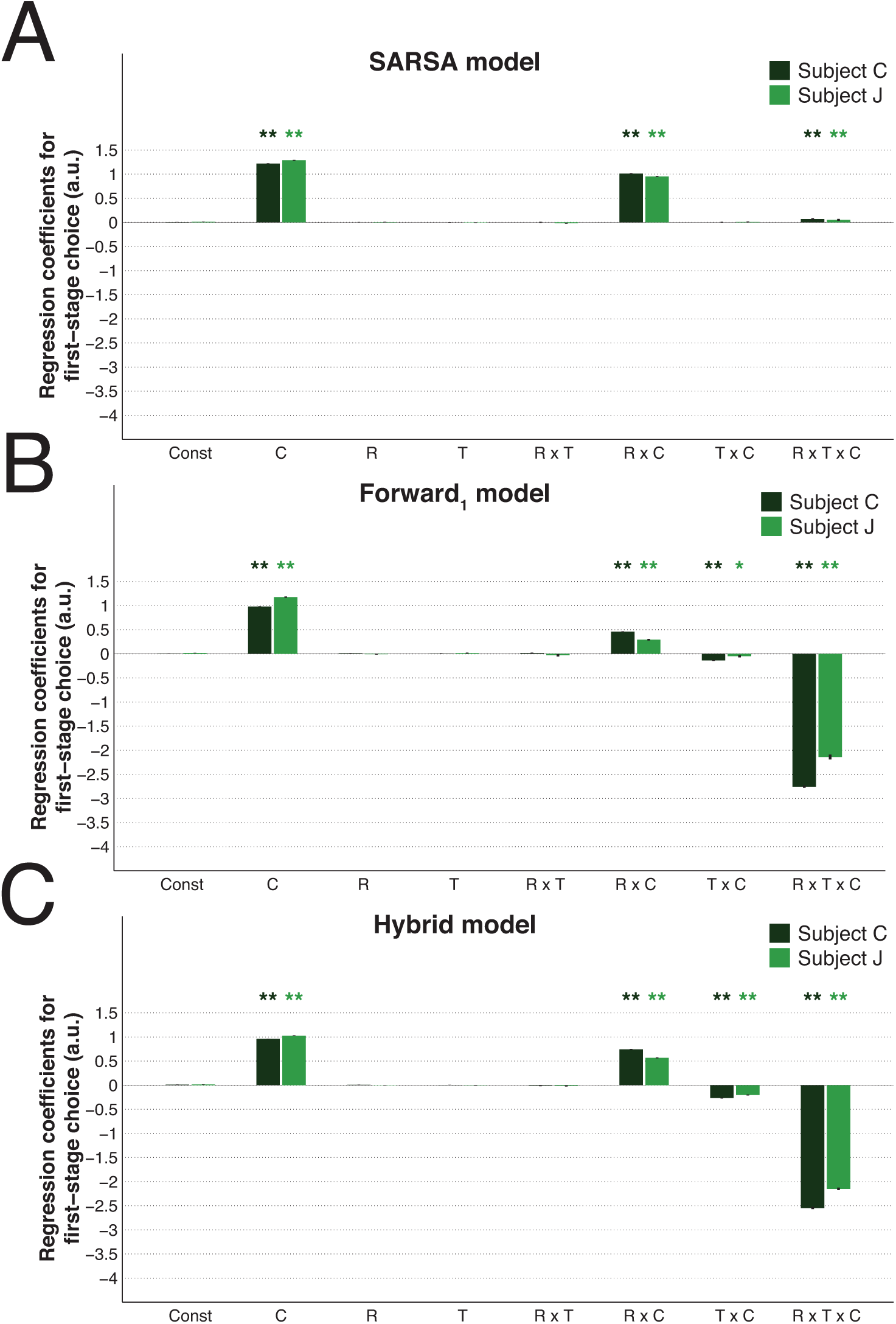
Graphical representation of the results from the logistic regression on first-stage simulated behaviour from each learning strategy, using the results from the previous trial’s predictor variables. The predictors used were: Const (constant term) captured any potential first-stage picture bias; C (previous first-stage choice; 1=car picture, 0=watering can picture) modelled a potential independent tendency to stick with the same option from trial to trial; R (previous outcome level; assumed as continuous and with low=1, medium=2, high=3), T (previous transition; rare=1, common=0) and R × T, measured any potential preference in first-stage picture choice given the previous outcome level, the previous transition and the interaction effect of both, respectively; R × C, T × C and R × T × C are the predictors of interest and quantify the main effects of reward, transition and the reward × transition interaction effect, respectively. All predictors were mean centred and continuous variables were also scaled by dividing them by two standard deviations (adjustments made before the computation of the interaction terms). Results for simulated choice behaviour (100 simulations per session for each subject and respecting the exact same reward structure) generated using the best-fitted mixed-effects parameters of the pure model-free *SARSA* model (A), pure model-sensitive *Forward*_1_ model (B) and *Hybrid* model (C). To note that the *Hybrid* model results are much closer to the MS-RL simulations as simulations used the parameters best fit to the subjects’ data and the MS weight estimated was close to 90%. Bar and error bar values correspond, respectively, to the mean and SE of the fixed-effects coefficients. ** for *α* = 0.01 and * for *α* = 0.05 in two-tailed one sample t-test with null-hypothesis mean equal to zero for the fixed-effects coefficients.

**Fig. S3.**
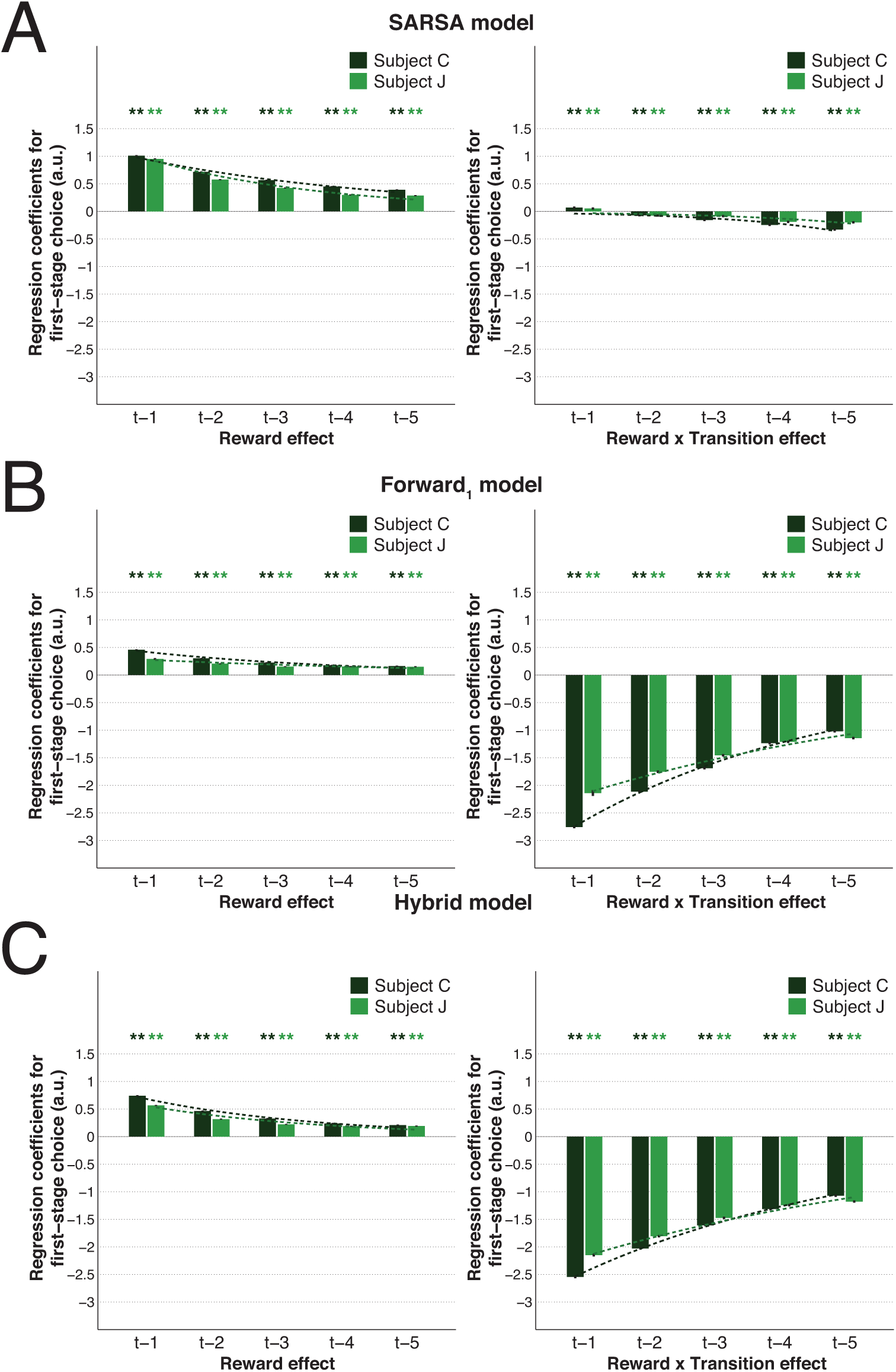
The impact of both reward and transition information from the five previous trials on first-stage simulated behaviour from each learning strategy. Multiple logistic regression results on first-stage simulated choice data (100 simulations per session for each subject and respecting the exact same reward structure) generated using the best-fitted mixed-effects parameters of the pure model-free *SARSA* model (A), pure model-sensitive *Forward*_1_ model (B) and *Hybrid* model (C) for the main effect of reward (left column) and reward × transition interaction term (right column) from the five previous trials. Bar and error bar values correspond, respectively, to the mean and SE of the fixed-effects coefficients. Dashed lines illustrate the exponential best fit on the mean fixed-effects coefficients of each trial into the past. ** for *α* = 0.01 and * for *α* = 0.05 in two-tailed one sample t-test with null-hypothesis mean equal to zero for the fixed-effects estimates.

**Fig. S4.**
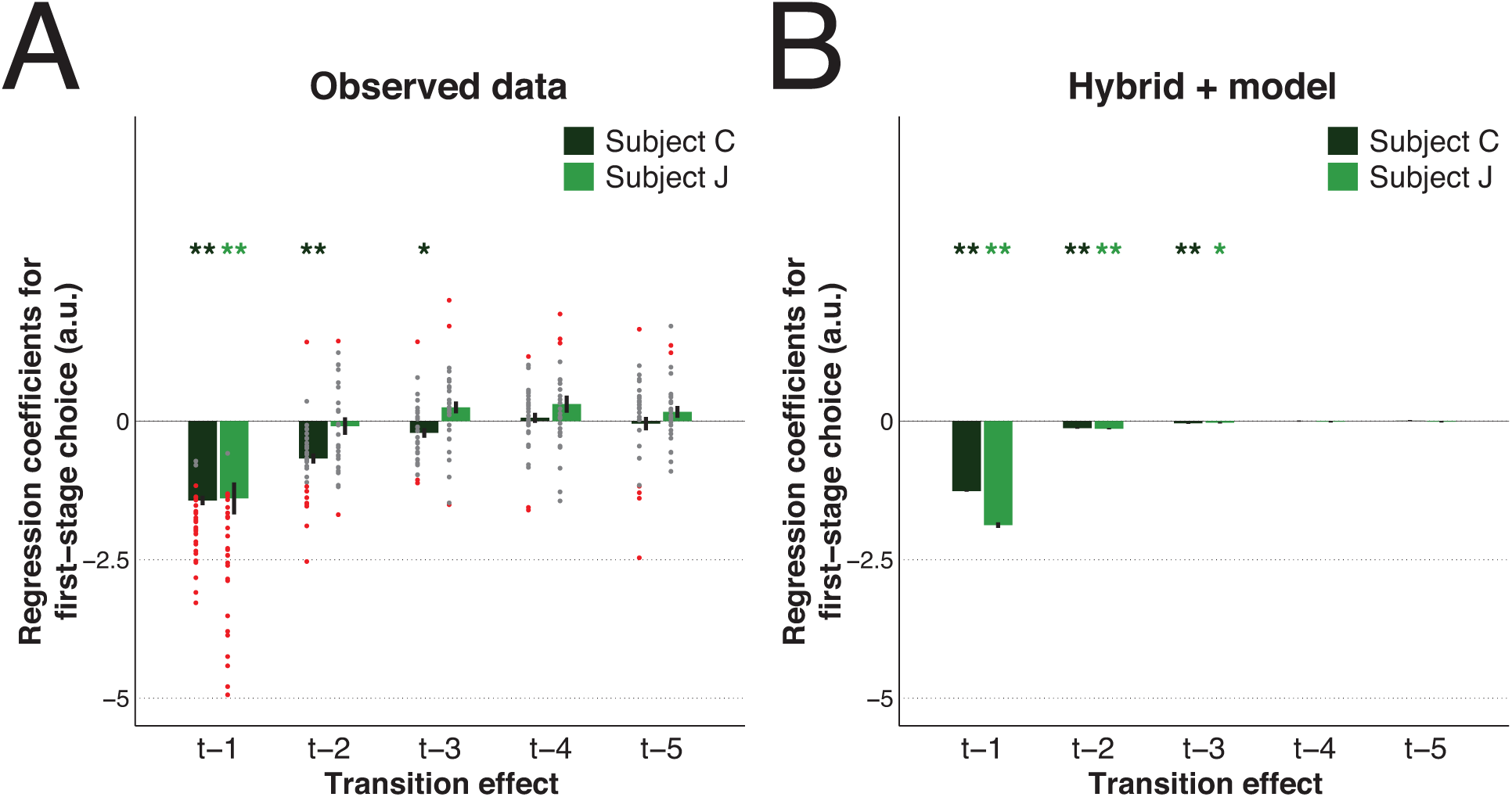
The impact of transition information on first-stage observed and simulated behaviour. Results of the main effect of transition from the five previous trials obtained in the logistic regression on observed first-stage choice (A) and on first-stage simulated choice data (B) generated using the best-fitted mixed-effects parameters of the *Hybrid*+ model (100 simulations per session for each subject and respecting the exact same reward structure). Dots represent the fixed-effects coefficients for each session (coloured red when *p* < 0.05 and grey otherwise). Bar and error bar values correspond, respectively, to the mixed-effect coefficients and their SE. ** *α* = 0.01 and * *α* = 0.05 in two-tailed one sample t-test with null-hypothesis mean equal to zero for the fixed-effects coefficients.

**Fig. S5.**
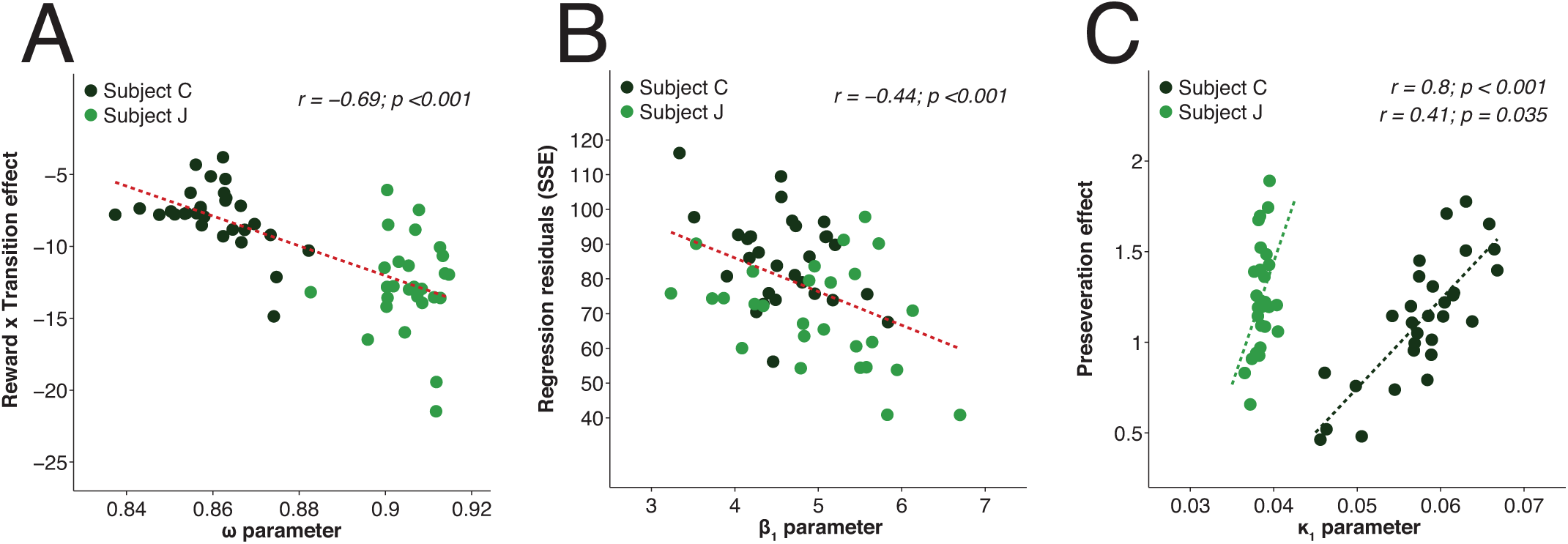
Correlation between logistic regression estimates and computational modelling parameters across sessions. (A) The greater the model-sensitive weight parameter *ω* obtained from the *Hybrid*+ model fitting, the more negative (i.e. the stronger the effect in the logistic regression) the regression coefficient for the reward × transition interaction (simulated results: *r* = −0.18, *p* < 0.001). (B) Relationship between the inverse temperature parameter at first-stage choice *β*_1_ obtained from the *Hybrid*+ model fitting and the residual values from the regression model (the greater the *β*_1_ parameter, the better the logistic regression fit; simulated results: *r* = −0.41, *p* < 0.001). (C) Positive correlation between the computational preseveration *κ*_1_ parameter and the regression coefficient for repeat first-stage choice independently of reward and transition (separate analysis for each subject because of the different *κ* parameters; simulated results: *r* = 0.35/0.08, *p* < 0.001/< 0.001 for C/J). Dashed lines represent the regression line of the fit for each individual subject or across subjects (in red). *r* is the Pearson’s linear correlation coefficients and *p* is the p-values across subjects in (A and B) and in (C) top values are for subject C and bottom values are for subject J.

**Fig. S6.**
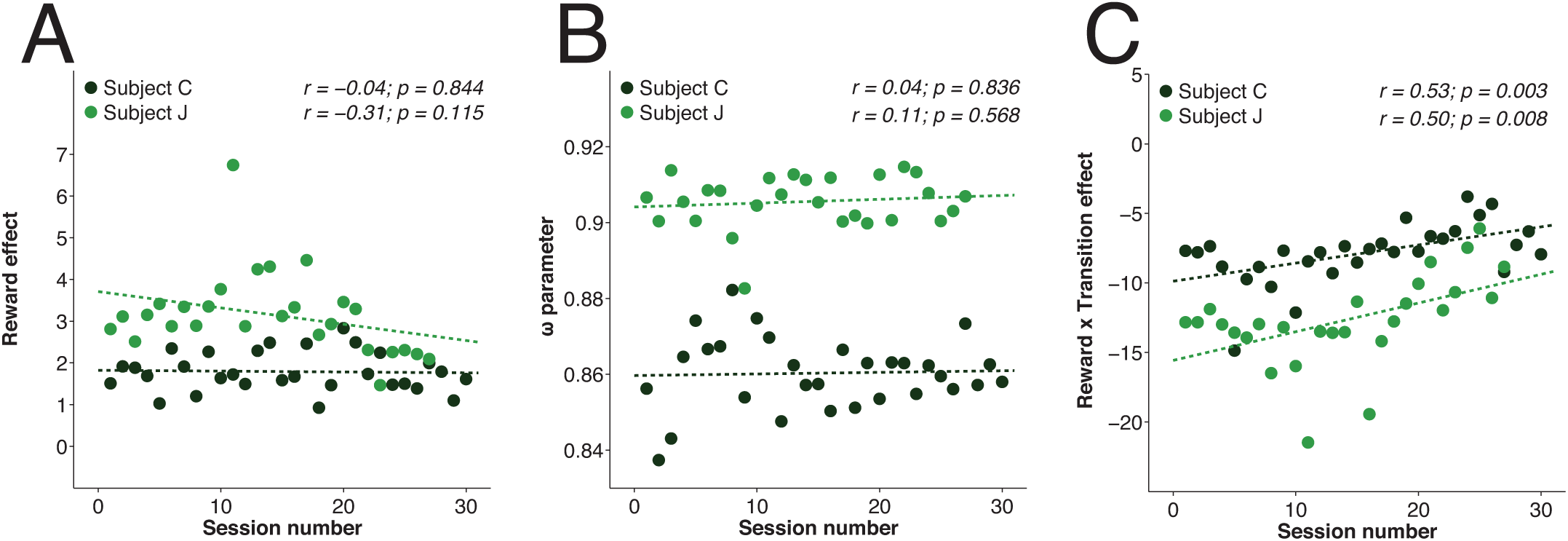
Evolution across sessions of logistic regression and computational modelling estimates. Across time and for both subjects, no significant decrease in the regression coefficients for the reward effect (A) or model-sensitive weight parameter *ω* (B) was found (both simulated results also with *p* > 0.05). However, a significant reduction was found for the effect of the regression coefficients for the reward × transition effect (C) with time (note that the more positive the regression coefficient the weaker the effect; simulated results: *r* = −0.01/−0.24, *p* = 0.959/0.228 for C/J). Dashed lines represent the regression line of the fit for each individual subject. *r* is the Pearson’s linear correlation coefficients and *p* is the p-values; top values are for subject C and bottom values are for subject J.

**Fig. S7.**
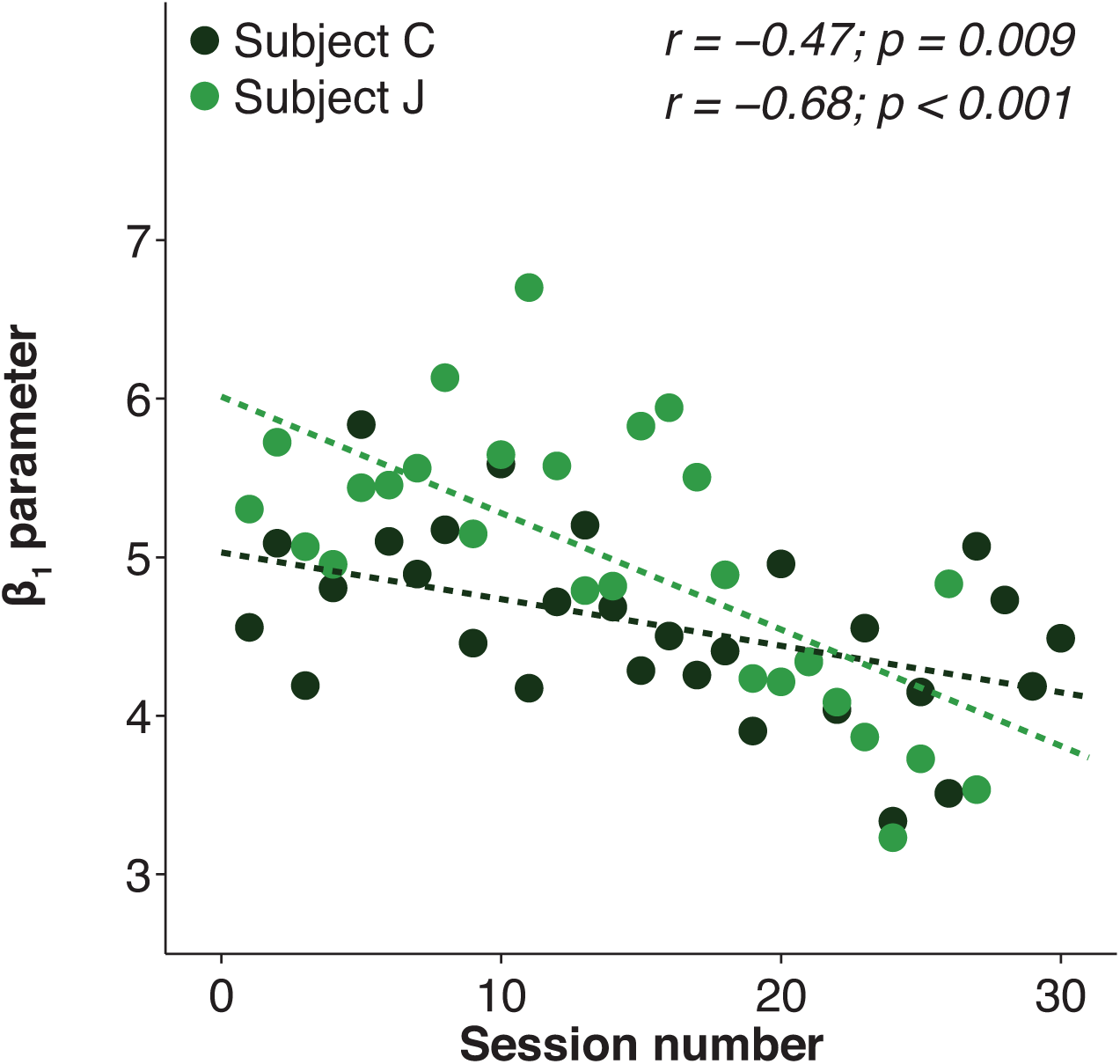
Evolution across sessions of the inverse temperature parameter for first-stage choice. As the number of sessions performed increased, subjects got progressively more stochastic (smaller inverse temperature values in observed behaviour; simulated results did not present such decrement: *r* = −0.02/−0.06, *p* = 0.898/0.768) in their choice behaviour. Dashed lines represent the regression line of the fit for each individual subject. *r* is the Pearson’s linear correlation coefficients and *p* is the p-values; top values are for subject C and bottom values are for subject J.

**Fig. S8.**
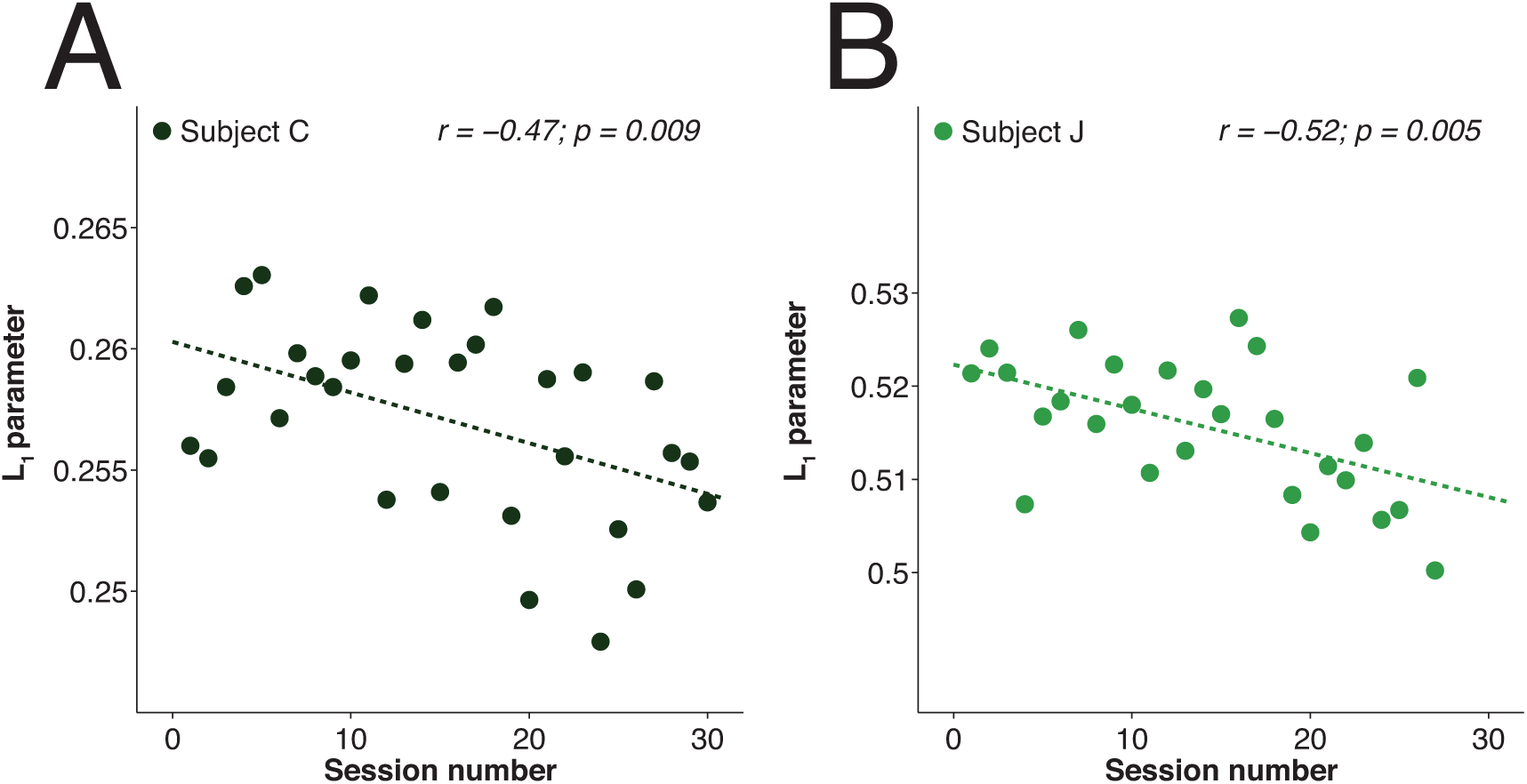
Evolution across sessions of the L_1_ parameter. As the number of sessions performed increased, the L_1_ parameter value got progressively smaller (i.e., less strength of the reinforcement by previous trial’s high reward) in both subjects. Dashed lines represent the regression line of the fit for each individual subject. *r* is the Pearson’s linear correlation coefficients and *p* is the p-values.

**Fig. S9.**
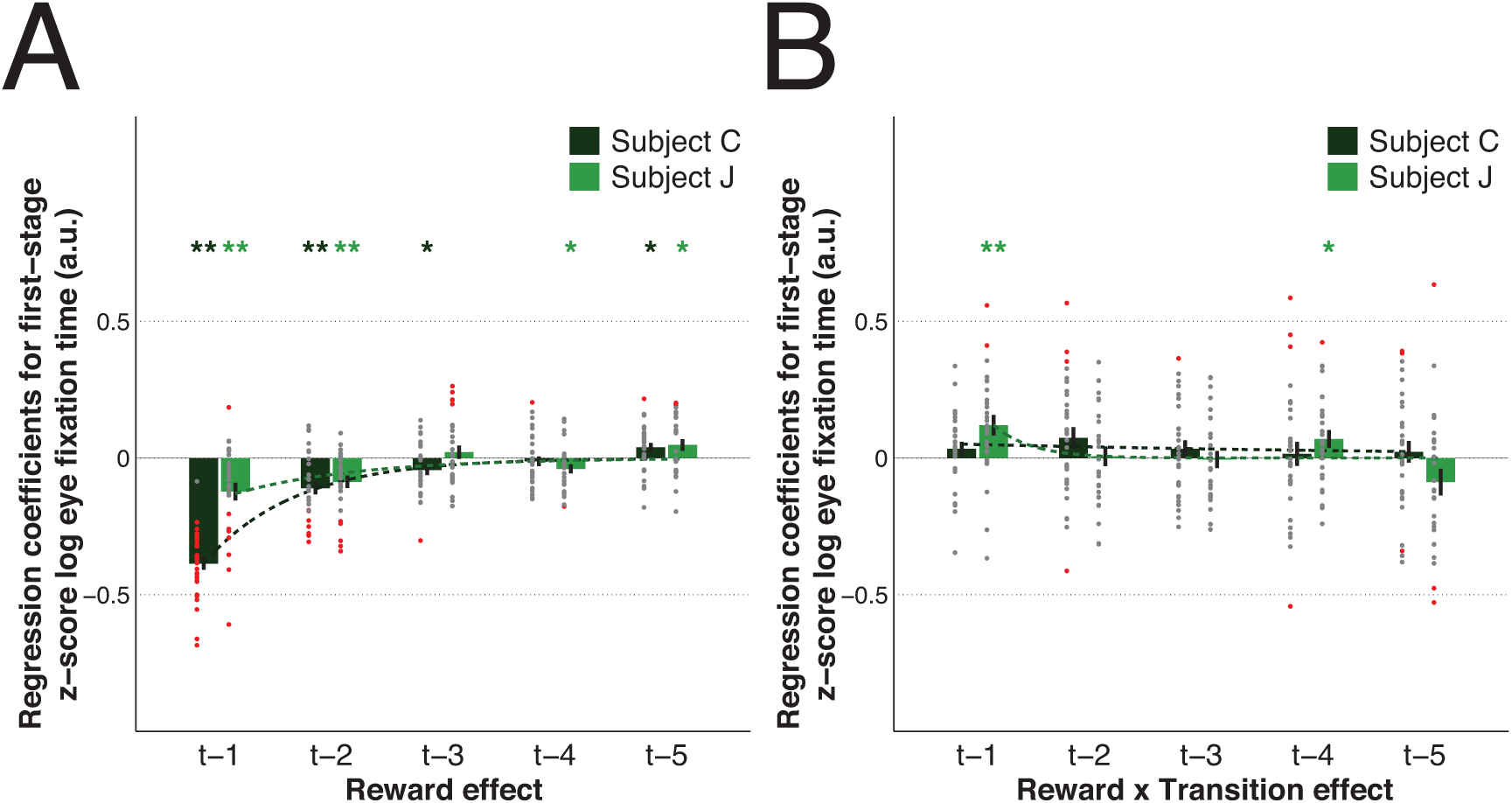
The impact of both reward and transition information on the first attempt to eye fixation at first-stage. Multiple linear regression results on *z*-scores of log transformed first-stage eye fixation time (high *z*-scores indicate slow first eye fixation attempt) with the contributions of the reward main effect (A) and reward × transition interaction term (B) from the five previous trials. Dots represent the fixed-effects coefficients for each session (coloured red when p < 0.05 and grey otherwise). Bar and error bar values correspond, respectively, to the mean value of the fixed-effect coefficients and its SEM. Dashed lines illustrate the exponential best fit on the mean fixed-effects coefficients of each trial into the past. ** for *α* = 0.01 and * for *α* = 0.05 in two-tailed one sample t-test with null-hypothesis mean equal to zero for the fixed-effects coefficients.

**Table S1.** Multiple logistic regression results for predictors of first-stage choice up to five trials back.

| Predictors<sup>‡</sup> | Fixed-effects* |  | Mixed-effects<sup>†</sup> |  |
| --- | --- | --- | --- | --- |
|  | C | J | C | J |
| Const | -0.11 (0.06) | -0.08 (0.08) | -0.06 (0.05) | -0.07 (0.08) |
| $C_{t-1}$ | 0.71 (0.06) | 0.91 (0.07)<sup>§</sup> | 0.56 (0.05)<sup>§</sup> | 0.57 (0.07)<sup>§</sup> |
| $R_{t-1}$ | -0.35 (0.06)<sup>§</sup> | -0.32 (0.11)<sup>§</sup> | -0.22 (0.06)<sup>§</sup> | -0.12 (0.22) |
| $T_{t-1}$ | 0.07 (0.05) | 0.07 (0.08) | 0.01 (0.05) | 0.05 (0.09) |
| $R_{t-1} \times T_{t-1}$ | 0.27 (0.12) | -0.18 (0.18) | 0.21 (0.11) | -0.45 (0.18)<sup>¶</sup> |
| $R_{t-1} \times C_{t-1}$ | *1.54 (0.10)<sup>§</sup>* | *3.26 (0.22)<sup>§</sup>* | *1.36 (0.09)<sup>§</sup>* | *2.82 (0.26)<sup>§</sup>* |
| $T_{t-1} \times C_{t-1}$ | -1.92 (0.11)<sup>§</sup> | -2.53 (0.21)<sup>§</sup> | -1.43 (0.08)<sup>§</sup> | -1.39 (0.29)<sup>§</sup> |
| $R_{t-1} \times T_{t-1} \times C_{t-1}$ | -7.85 (0.50)<sup>§</sup> | -13.76 (0.70)<sup>§</sup> | -7.06 (0.39)<sup>§</sup> | -16.37 (1.22)<sup>§</sup> |
| $C_{t-2}$ | 0.37 (0.05)<sup>§</sup> | 0.17 (0.08) | 0.32 (0.04)<sup>§</sup> | 0.11 (0.07) |
| $R_{t-2}$ | -0.05 (0.05) | 0.03 (0.08) | -0.03 (0.05) | 0.03 (0.08) |
| $T_{t-2}$ | 0.11 (0.06) | 0.07 (0.07) | 0.06 (0.04) | 0.08 (0.05) |
| $R_{t-2} \times T_{t-2}$ | 0.28 (0.11)<sup>¶</sup> | 0.17 (0.14) | 0.14 (0.09) | 0.14 (0.14) |
| $R_{t-2} \times C_{t-2}$ | *0.70 (0.10)<sup>§</sup>* | *0.57 (0.12)<sup>§</sup>* | *0.62 (0.08)<sup>§</sup>* | *0.63 (0.13)<sup>§</sup>* |
| $T_{t-2} \times C_{t-2}$ | -0.72 (0.13)<sup>§</sup> | -0.13 (0.15) | -0.67 (0.09)<sup>§</sup> | -0.09 (0.16) |
| $R_{t-2} \times T_{t-2} \times C_{t-2}$ | -2.75 (0.31)<sup>§</sup> | -2.68 (0.33)<sup>§</sup> | -2.43 (0.26)<sup>§</sup> | -2.11 (0.32)<sup>§</sup> |
| $C_{t-3}$ | 0.17 (0.06)<sup>§</sup> | 0.07 (0.09) | 0.17 (0.05)<sup>§</sup> | 0.06 (0.08) |
| $R_{t-3}$ | 0.12 (0.06) | -0.02 (0.08) | 0.07 (0.04) | -0.03 (0.08) |
| $T_{t-3}$ | 0.15 (0.06)<sup>¶</sup> | 0.07 (0.06) | 0.12 (0.04)<sup>§</sup> | 0.09 (0.05) |
| $R_{t-3} \times T_{t-3}$ | 0.09 (0.11) | -0.19 (0.15) | 0.11 (0.10) | -0.14 (0.15) |
| $R_{t-3} \times C_{t-3}$ | *0.31 (0.11)<sup>§</sup>* | *0.30 (0.17)* | *0.33 (0.08)<sup>§</sup>* | *0.28 (0.13)<sup>¶</sup>* |
| $T_{t-3} \times C_{t-3}$ | -0.26 (0.11)<sup>¶</sup> | *0.19 (0.15)* | -0.21 (0.09)<sup>¶</sup> | *0.25 (0.11)<sup>¶</sup>* |
| $R_{t-3} \times T_{t-3} \times C_{t-3}$ | -1.33 (0.22)<sup>§</sup> | -1.31 (0.40)<sup>§</sup> | -1.26 (0.19)<sup>§</sup> | -1.48 (0.32)<sup>§</sup> |
| $C_{t-4}$ | 0.05 (0.06) | -0.07 (0.07) | 0.04 (0.04) | -0.05 (0.06) |
| $R_{t-4}$ | 0.03 (0.06) | -0.04 (0.06) | -0.19 (0.06) | -0.04 (0.05) |
| $T_{t-4}$ | 0.02 (0.06) | 0.04 (0.06) | 0.02 (0.06) | 0.03 (0.05) |
| $R_{t-4} \times T_{t-4}$ | 0.04 (0.10) | -0.11 (0.15) | 0.06 (0.09) | -0.17 (0.17) |
| $R_{t-4} \times C_{t-4}$ | *0.23 (0.10)<sup>¶</sup>* | *0.07 (0.11)* | *0.20 (0.08)<sup>¶</sup>* | *0.17 (0.11)* |
| $T_{t-4} \times C_{t-4}$ | *0.06 (0.12)* | *0.23 (0.14)* | *0.06 (0.09)* | *0.31 (0.15)<sup>¶</sup>* |
| $R_{t-4} \times T_{t-4} \times C_{t-4}$ | -0.70 (0.25)<sup>§</sup> | -0.89 (0.29)<sup>§</sup> | -0.66 (0.19)<sup>§</sup> | -1.02 (0.32)<sup>§</sup> |
| $C_{t-5}$ | 0.06 (0.05) | 0.15 (0.05)<sup>¶</sup> | 0.06 (0.04) | 0.11 (0.05)<sup>¶</sup> |
| $R_{t-5}$ | 0.10 (0.05) | -0.02 (0.05) | 0.11 (0.04)<sup>§</sup> | -0.04 (0.06) |
| $T_{t-5}$ | 0.07 (0.05) | 0.01 (0.05) | 0.02 (0.04) | 0.02 (0.05) |
| $R_{t-5} \times T_{t-5}$ | 0.05 (0.10) | -0.05 (0.16) | 0.08 (0.09) | 0.12 (0.15) |
| $R_{t-5} \times C_{t-5}$ | *0.01 (0.12)* | *0.10 (0.10)* | -0.01 (0.11) | *0.01 (0.10)* |
| $T_{t-5} \times C_{t-5}$ | -0.02 (0.15) | *0.22 (0.11)* | -0.04 (0.12) | *0.17 (0.11)* |
| $R_{t-5} \times T_{t-5} \times C_{t-5}$ | -0.50 (0.24)<sup>¶</sup> | *0.22 (0.26)* | -0.35 (0.22) | -0.34 (0.29) |

\* Values of fixed-effects results are mean (SEM) of the regression coefficients across sessions.
<sup>†</sup> Values of mixed-effects results are the regression coefficients (SE).
<sup>‡</sup> For the given trial $t$ , the variables used were: The predictors used were: Const (constant term) captured any potential first-stage picture bias; C (previous first-stage choice; 1=car picture, 0=watering can picture) modelled a potential independent tendency to stick with the same option from trial to trial; R (previous outcome level; assumed as continuous and with low=1, medium=2, high=3), T (previous transition; rare=1, common=0) and $R \times T$ , measured any potential preference in first-stage picture choice given the previous outcome level, the previous transition and the interaction effect of both, respectively; $R \times C$ , $T \times C$ and $R \times T \times C$ are the predictors of interest (in italic) and quantify the main effects of reward, transition and the reward × transition interaction effect, respectively. All predictors were mean centred and continuous variables were also scaled by dividing them by two SD (adjustments made before the computation of the interaction terms).
<sup>§</sup> Significance at the 0.01 level. <sup>¶</sup> Significance at the 0.05 level.

**Table S2.** Model comparison results for the model-free *SARSA* models.

| Parameters* | Fixed-effects *BIC* sum |  | Mixed-effects *BIC*<sub>int</sub> |  |
| --- | --- | --- | --- | --- |
|  | C | J | C | J |
| $\alpha, \beta, \kappa_1$ | 35873 | 34670 | 35717 | 34538 |
| $\alpha, \beta, \kappa_2$ | 36581 | 35191 | 36444 | 35170 |
| $\alpha, \beta, \kappa$ | 35814 | 34133 | 35679 | 34043 |
| $\alpha, \beta, \lambda$ | 36330 | 35157 | 36149 | 35171 |
| $\alpha_1, \alpha_2, \beta$ | 36511 | 35671 | 36494 | 35700 |
| $\alpha, \beta_1, \beta_2$ | 36532 | 35682 | 36393 | 35596 |
| $\alpha_1, \alpha_2, \beta_1, \beta_2$ | 36615 | 35770 | 36464 | 35775 |
| $\alpha_1, \alpha_2, \beta, \kappa_1$ | 35822 | 34759 | 35689 | 34588 |
| $\alpha_1, \alpha_2, \beta, \kappa_2$ | 36584 | 35295 | 36430 | 35274 |
| $\alpha_1, \alpha_2, \beta, \kappa$ | 35821 | 34252 | 35687 | 34087 |
| $\alpha_1, \alpha_2, \beta, \lambda$ | 36335 | 35267 | 36116 | 35231 |
| $\alpha, \beta_1, \beta_2, \kappa_1$ | 35785 | 34742 | 35455 | 34458 |
| $\alpha, \beta_1, \beta_2, \kappa_2$ | 36619 | 35306 | 36348 | 35175 |
| $\alpha, \beta_1, \beta_2, \kappa$ | 35934 | 34267 | 35637 | 34022 |
| $\alpha, \beta_1, \beta_2, \lambda$ | 36447 | 35260 | 36163 | 35176 |
| $\alpha, \beta, \kappa_1, \kappa_2$ | 35908 | 34256 | 35624 | 34069 |
| $\alpha, \beta, \kappa_1, \lambda$ | 35723 | 34415 | 35424 | 34185 |
| $\alpha, \beta, \kappa_2, \lambda$ | 36403 | 34825 | 36095 | 34791 |
| $\alpha, \beta, \kappa, \lambda$ | **35697<sup>†</sup>** | **33931<sup>†</sup>** | 35420 | **33750<sup>†</sup>** |
| $\alpha_1, \alpha_2, \beta_1, \beta_2, \kappa_1$ | 35895 | 34859 | 35494 | 34509 |
| $\alpha_1, \alpha_2, \beta_1, \beta_2, \kappa_2$ | 37797 | 35690 | 36765 | 35698 |
| $\alpha_1, \alpha_2, \beta_1, \beta_2, \kappa$ | 35949 | 34388 | 35723 | 34088 |
| $\alpha_1, \alpha_2, \beta_1, \beta_2, \lambda$ | 36478 | 35360 | 36142 | 35272 |
| $\alpha, \beta_1, \beta_2, \kappa_1, \kappa_2$ | 35873 | 34370 | 35408 | 34049 |
| $\alpha, \beta_1, \beta_2, \kappa_1, \lambda$ | 35794 | 34553 | 35319 | 34183 |
| $\alpha, \beta_1, \beta_2, \kappa_2, \lambda$ | 36531 | 34888 | 36117 | 34774 |
| $\alpha, \beta_1, \beta_2, \kappa, \lambda$ | 35856 | 34058 | 35385 | 33754 |
| $\alpha_1, \alpha_2, \beta, \kappa_1, \kappa_2$ | 35891 | 34373 | 36074 | 34133 |
| $\alpha_1, \alpha_2, \beta, \kappa_1, \lambda$ | 35707 | 34539 | 35356 | 34240 |
| $\alpha_1, \alpha_2, \beta, \kappa_2, \lambda$ | 36426 | 34936 | 36074 | 34856 |
| $\alpha_1, \alpha_2, \beta, \kappa, \lambda$ | 35725 | 34061 | 35382 | 33788 |
| $\alpha, \beta, \kappa_1, \kappa_2, \lambda$ | 35784 | 34055 | 35348 | 33780 |
| $\alpha_1, \alpha_2, \beta_1, \beta_2, \kappa_1, \kappa_2$ | 35983 | 34494 | 35439 | 34087 |
| $\alpha_1, \alpha_2, \beta_1, \beta_2, \kappa_1, \lambda$ | 35858 | 34677 | 35311 | 34233 |
| $\alpha_1, \alpha_2, \beta_1, \beta_2, \kappa_2, \lambda$ | 36566 | 34994 | 36090 | 34865 |
| $\alpha_1, \alpha_2, \beta_1, \beta_2, \kappa, \lambda$ | 35858 | 34188 | 35331 | 33796 |
| $\alpha, \beta_1, \beta_2, \kappa_1, \kappa_2, \lambda$ | 35880 | 34186 | 35266 | 33766 |
| $\alpha_1, \alpha_2, \beta, \kappa_1, \kappa_2, \lambda$ | 35791 | 34186 | 35287 | 33824 |
| $\alpha_1, \alpha_2, \beta_1, \beta_2, \kappa_1, \kappa_2, \lambda$ | 35946 | 34312 | **35260<sup>†</sup>** | 33825 |

\* Abbreviations: learning rate for first-stage ( $\alpha_1$ ) and second-stage ( $\alpha_2$ ); $\alpha$ is when $\alpha_1 = \alpha_2$ ; inverse temperature for first-stage ( $\beta_1$ ) and second-stage ( $\beta_2$ ); $\beta$ is when $\beta_1 = \beta_2$ ; perseveration for first-stage ( $\kappa_1$ ) and second-stage ( $\kappa_2$ ); $\kappa$ is when $\kappa_1 = \kappa_2$ ; eligibility trace ( $\lambda$ ).
<sup>†</sup> Best fitting *SARSA* model variant for the respective subject and analysis type.

**Table S3.** Model comparison results for the model-free *Q*-learning models.

| Parameters * | Fixed-effects $BIC$ sum | | Mixed-effects $BIC_{int}$ | |
| --- | --- | --- | --- | --- |
|  | C | J | C | J |
| $\alpha, \beta$ | 36445 | 35620 | 36415 | 35706 |
| $\alpha, \beta, \kappa_1$ | 35767 | 34668 | 35633 | 34574 |
| $\alpha, \beta, \kappa_2$ | 36508 | 35252 | 36354 | 35283 |
| $\alpha, \beta, \kappa$ | 35736 | 34162 | 35620 | 34101 |
| $\alpha, \beta, \lambda$ | 36357 | 35399 | 36245 | 35525 |
| $\alpha_1, \alpha_2, \beta$ | 36484 | 35729 | 36428 | 35749 |
| $\alpha, \beta_1, \beta_2$ | 36501 | 35731 | 36408 | 35764 |
| $\alpha_1, \alpha_2, \beta_1, \beta_2$ | 36609 | 35836 | 36431 | 35878 |
| $\alpha_1, \alpha_2, \beta, \kappa_1$ | 35778 | 34788 | 35614 | 34607 |
| $\alpha_1, \alpha_2, \beta, \kappa_2$ | 36566 | 35372 | 36373 | 35427 |
| $\alpha_1, \alpha_2, \beta, \kappa$ | 35792 | 34297 | 35615 | 34088 |
| $\alpha_1, \alpha_2, \beta, \lambda$ | 36420 | 35502 | 36265 | 35556 |
| $\alpha, \beta_1, \beta_2, \kappa_1$ | 35761 | 34780 | 35483 | 34589 |
| $\alpha, \beta_1, \beta_2, \kappa_2$ | 36587 | 35357 | 36370 | 35383 |
| $\alpha, \beta_1, \beta_2, \kappa$ | 35878 | 34300 | 35609 | 34084 |
| $\alpha, \beta_1, \beta_2, \lambda$ | 36458 | 35494 | 36367 | 35528 |
| $\alpha, \beta, \kappa_1, \kappa_2$ | 35819 | 34281 | 35560 | 34127 |
| $\alpha, \beta, \kappa_1, \lambda$ | 35721 | 34571 | 35486 | 34470 |
| $\alpha, \beta, \kappa_2, \lambda$ | 36434 | 35068 | 36206 | 35160 |
| $\alpha, \beta, \kappa, \lambda$ | **35709<sup>†</sup>** | **34095<sup>†</sup>** | 35502 | 33942 |
| $\alpha_1, \alpha_2, \beta_1, \beta_2, \kappa_1$ | 35891 | 34909 | 35559 | 34545 |
| $\alpha_1, \alpha_2, \beta_1, \beta_2, \kappa_2$ | 37797 | 35773 | 36742 | 35552 |
| $\alpha_1, \alpha_2, \beta_1, \beta_2, \kappa$ | 35925 | 34426 | 35829 | 34119 |
| $\alpha_1, \alpha_2, \beta_1, \beta_2, \lambda$ | 36543 | 35590 | 36312 | 35612 |
| $\alpha, \beta_1, \beta_2, \kappa_1, \kappa_2$ | 35849 | 34409 | 35451 | 34140 |
| $\alpha, \beta_1, \beta_2, \kappa_1, \lambda$ | 35799 | 34713 | 35411 | 34352 |
| $\alpha, \beta_1, \beta_2, \kappa_2, \lambda$ | 36540 | 35121 | 36301 | 35121 |
| $\alpha, \beta_1, \beta_2, \kappa, \lambda$ | 35856 | 34220 | 35467 | **33935<sup>†</sup>** |
| $\alpha_1, \alpha_2, \beta, \kappa_1, \kappa_2$ | 35856 | 34413 | 35537 | 34169 |
| $\alpha_1, \alpha_2, \beta, \kappa_1, \lambda$ | 35762 | 34704 | 35445 | 34423 |
| $\alpha_1, \alpha_2, \beta, \kappa_2, \lambda$ | 36510 | 35143 | 36227 | 35169 |
| $\alpha_1, \alpha_2, \beta, \kappa, \lambda$ | 35786 | 34224 | 35492 | 33969 |
| $\alpha, \beta, \kappa_1, \kappa_2, \lambda$ | 35788 | 34217 | 35437 | 34006 |
| $\alpha_1, \alpha_2, \beta_1, \beta_2, \kappa_1, \kappa_2$ | 35979 | 34546 | 36026 | 34133 |
| $\alpha_1, \alpha_2, \beta_1, \beta_2, \kappa_1, \lambda$ | 35904 | 34844 | 35399 | 34432 |
| $\alpha_1, \alpha_2, \beta_1, \beta_2, \kappa_2, \lambda$ | 36566 | 34994 | 36084 | 34888 |
| $\alpha_1, \alpha_2, \beta_1, \beta_2, \kappa, \lambda$ | 35907 | 34350 | 35524 | 33990 |
| $\alpha, \beta_1, \beta_2, \kappa_1, \kappa_2, \lambda$ | 35884 | 34347 | 35429 | 34010 |
| $\alpha_1, \alpha_2, \beta, \kappa_1, \kappa_2, \lambda$ | 35847 | 34346 | 35404 | 34038 |
| $\alpha_1, \alpha_2, \beta_1, \beta_2, \kappa_1, \kappa_2, \lambda$ | 35992 | 34473 | **35376<sup>†</sup>** | 34008 |

\* Abbreviations: learning rate for first-stage ( $\alpha_1$ ) and second-stage ( $\alpha_2$ ); $\alpha$ is when $\alpha_1 = \alpha_2$ ; inverse temperature for first-stage ( $\beta_1$ ) and second-stage ( $\beta_2$ ); $\beta$ is when $\beta_1 = \beta_2$ ; perseveration for first-stage ( $\kappa_1$ ) and second-stage ( $\kappa_2$ ); $\kappa$ is when $\kappa_1 = \kappa_2$ ; eligibility trace ( $\lambda$ ).
<sup>†</sup> Best fitting $Q$ -learning model variant for the respective subject and analysis type.

**Table S4.** Model comparison results for the three model-sensitive models.

| Model* | Parameters<sup>†</sup> | Fixed-effects $BIC$ sum | | Mixed-effects $BIC_{int}$ | |
| --- | --- | --- | --- | --- | --- |
|  |  | C | J | C | J |
| $Forward_1$ | | | | | |
| | $\alpha_2, \beta$ | 35298 | 34824 | 35275 | 34868 |
| | $\alpha_2, \beta_1, \beta_2$ | 34630 | 33708 | 34548 | 33743 |
| | $\alpha_2, \beta, \kappa_1$ | 34737 | 34038 | 34610 | 33965 |
| | $\alpha_2, \beta, \kappa_2$ | 35425 | 34572 | 35271 | 34619 |
| | $\alpha_2, \beta, \kappa$ | 34818 | 33634 | 34701 | 33616 |
| | $\alpha_2, \beta, \kappa_1, \kappa_2$ | 34856 | 33753 | 34595 | 33658 |
| | $\alpha_2, \beta_1, \beta_2, \kappa_1$ | 34418 | 33437 | 34176 | 33345 |
| | $\alpha_2, \beta_1, \beta_2, \kappa_2$ | 34715 | 33342 | 34499 | 33309 |
| | $\alpha_2, \beta_1, \beta_2, \kappa$ | **34360<sup>‡</sup>** | 33182 | **34122<sup>‡</sup>** | 33248 |
| | $\alpha_2, \beta_1, \beta_2, \kappa_1, \kappa_2$ | 34505 | **33071<sup>‡</sup>** | 34143 | **32837<sup>‡</sup>** |
| $Forward_2$ | | | | | |
| | $\alpha_2, \beta$ | 35304 | 34831 | 35279 | 34872 |
| | $\alpha_2, \beta_1, \beta_2$ | 34642 | 33732 | 34556 | 33758 |
| | $\alpha_2, \beta, \kappa_1$ | 34743 | 34046 | 34610 | 33959 |
| | $\alpha_2, \beta, \kappa_2$ | 35432 | 34579 | 35274 | 34612 |
| | $\alpha_2, \beta, \kappa$ | 34824 | 33641 | 34702 | 33619 |
| | $\alpha_2, \beta, \kappa_1, \kappa_2$ | 34862 | 33761 | 34594 | 33661 |
| | $\alpha_2, \beta_1, \beta_2, \kappa_1$ | 34430 | 33462 | 34181 | 33359 |
| | $\alpha_2, \beta_1, \beta_2, \kappa_2$ | 34727 | 33366 | 34508 | 33324 |
| | $\alpha_2, \beta_1, \beta_2, \kappa$ | 34372 | 33205 | 34129 | 33256 |
| | $\alpha_2, \beta_1, \beta_2, \kappa_1, \kappa_2$ | 34517 | 33095 | 34152 | 32851 |
| $Forward_3$ | | | | | |
| | $\alpha_2, \beta, \zeta$ | 35484 | 34993 | 35297 | 34883 |
| | $\alpha_2, \beta_1, \beta_2, \zeta$ | 34797 | 33841 | 34570 | 33748 |
| | $\alpha_2, \beta, \kappa_1, \zeta$ | 34923 | 34206 | 34630 | 33965 |
| | $\alpha_2, \beta, \kappa_2, \zeta$ | 35611 | 34742 | 35288 | 34633 |
| | $\alpha_2, \beta, \kappa, \zeta$ | 35004 | 33803 | 34717 | 33629 |
| | $\alpha_2, \beta, \kappa_1, \kappa_2, \zeta$ | 35042 | 33922 | 34624 | 33656 |
| | $\alpha_2, \beta_1, \beta_2, \kappa_1, \zeta$ | 34590 | 33579 | 34192 | 33346 |
| | $\alpha_2, \beta_1, \beta_2, \kappa_2, \zeta$ | 34882 | 33475 | 34516 | 33313 |
| | $\alpha_2, \beta_1, \beta_2, \kappa, \zeta$ | 34533 | 33335 | 34140 | 33255 |
| | $\alpha_2, \beta_1, \beta_2, \kappa_1, \kappa_2, \zeta$ | 35229 | 34091 | 34145 | 32841 |

\*See *SI Methods* for main differences between each of the three model-sensitive (MS) models use.
<sup>†</sup>Abbreviations: learning rate for second-stage ( $\alpha_2$ ); inverse temperature for first-stage ( $\beta_1$ ) and second-stage ( $\beta_2$ ); $\beta$ is when $\beta_1 = \beta_2$ ; perseveration for first-stage ( $\kappa_1$ ) and second-stage ( $\kappa_2$ ); $\kappa$ is when $\kappa_1 = \kappa_2$ ; eligibility trace ( $\lambda$ ); $\zeta$ is a weight given to state-transition model testing.
<sup>‡</sup>Best fitting MS model variant for the respective subject and analysis type.

**Table S5.** Model comparison results for the *Hybrid* models.

| Parameters* | Fixed-effects $BIC$ sum | | Mixed-effects $BIC_{int}$ | |
| --- | --- | --- | --- | --- |
|  | C | J | C | J |
| $\alpha, \beta, \beta, \omega$ | 35306 | 34893 | 35148 | 34834 |
| $\alpha, \beta, \kappa_1, \omega$ | 34807 | 34144 | 34522 | 33930 |
| $\alpha, \beta, \kappa_2, \omega$ | 35435 | 34642 | 35148 | 34592 |
| $\alpha, \beta, \kappa, \omega$ | 34880 | 33742 | 34616 | 33600 |
| $\alpha, \beta, \lambda, \omega$ | 35451 | 34900 | 35171 | 34775 |
| $\alpha_1, \alpha_2, \beta, \omega$ | 35464 | 35028 | 35168 | 34814 |
| $\alpha, \beta_1, \beta_2, \omega$ | 34515 | 33638 | 34246 | 33548 |
| $\alpha_1, \alpha_2, \beta_1, \beta_2, \omega$ | 34577 | 33735 | 34313 | 33585 |
| $\alpha_1, \alpha_2, \beta, \kappa_1, \omega$ | 34973 | 34292 | 34561 | 33923 |
| $\alpha_1, \alpha_2, \beta, \kappa_2, \omega$ | 35593 | 34779 | 35195 | 34582 |
| $\alpha_1, \alpha_2, \beta, \kappa, \omega$ | 35045 | 33891 | 34644 | 33617 |
| $\alpha_1, \alpha_2, \beta, \lambda, \omega$ | 35602 | 35023 | 35191 | 34733 |
| $\alpha, \beta_1, \beta_2, \kappa_1, \omega$ | 34380 | 33432 | 33948 | 33215 |
| $\alpha, \beta_1, \beta_2, \kappa_2, \omega$ | 34602 | 33269 | 34194 | 33217 |
| $\alpha, \beta_1, \beta_2, \kappa, \omega$ | **34326<sup>†</sup>** | 33199 | **33898<sup>†</sup>** | 33189 |
| $\alpha, \beta_1, \beta_2, \lambda, \omega$ | 34652 | 33640 | 34244 | 33499 |
| $\alpha, \beta, \kappa_1, \kappa_2, \omega$ | 34928 | 33861 | 34508 | 33673 |
| $\alpha, \beta, \kappa_1, \lambda, \omega$ | 34966 | 34216 | 34542 | 33913 |
| $\alpha, \beta, \kappa_2, \lambda, \omega$ | 35580 | 34650 | 35167 | 34542 |
| $\alpha, \beta, \kappa, \lambda, \omega$ | 35036 | 33813 | 34640 | 33566 |
| $\alpha_1, \alpha_2, \beta_1, \beta_2, \kappa_1, \omega$ | 34468 | 33553 | 33986 | 33252 |
| $\alpha_1, \alpha_2, \beta_1, \beta_2, \kappa_2, \omega$ | 34663 | 33367 | 34265 | 33167 |
| $\alpha_1, \alpha_2, \beta_1, \beta_2, \kappa, \omega$ | 34422 | 33346 | 33948 | 33239 |
| $\alpha_1, \alpha_2, \beta_1, \beta_2, \lambda, \omega$ | 34709 | 33748 | 34302 | 33541 |
| $\alpha, \beta_1, \beta_2, \kappa_1, \kappa_2, \omega$ | 34468 | **33063<sup>†</sup>** | 33904 | **32807<sup>†</sup>** |
| $\alpha, \beta_1, \beta_2, \kappa_1, \lambda, \omega$ | 34528 | 33473 | 33952 | 33182 |
| $\alpha, \beta_1, \beta_2, \kappa_2, \lambda, \omega$ | 34739 | 33272 | 34202 | 33075 |
| $\alpha, \beta_1, \beta_2, \kappa, \lambda, \omega$ | 34475 | 33249 | 33906 | 33197 |
| $\alpha_1, \alpha_2, \beta, \kappa_1, \kappa_2, \omega$ | 35095 | 34010 | 34558 | 33642 |
| $\alpha_1, \alpha_2, \beta, \kappa_1, \lambda, \omega$ | 35125 | 34355 | 34576 | 33889 |
| $\alpha_1, \alpha_2, \beta, \kappa_2, \lambda, \omega$ | 35731 | 34769 | 35200 | 34552 |
| $\alpha_1, \alpha_2, \beta, \kappa, \lambda, \omega$ | 35194 | 33951 | 34668 | 33566 |
| $\alpha, \beta, \kappa_1, \kappa_2, \lambda, \omega$ | 35087 | 33935 | 34524 | 33612 |
| $\alpha_1, \alpha_2, \beta_1, \beta_2, \kappa_1, \kappa_2, \omega$ | 34556 | 33186 | 33950 | 32843 |
| $\alpha_1, \alpha_2, \beta_1, \beta_2, \kappa_1, \lambda, \omega$ | 34612 | 33599 | 33986 | 33234 |
| $\alpha_1, \alpha_2, \beta_1, \beta_2, \kappa_2, \lambda, \omega$ | 34795 | 33380 | 34255 | 33180 |
| $\alpha_1, \alpha_2, \beta_1, \beta_2, \kappa, \lambda, \omega$ | 34568 | 33392 | 33940 | 33235 |
| $\alpha, \beta_1, \beta_2, \kappa_1, \kappa_2, \lambda, \omega$ | 34616 | 33105 | 33903 | 32917 |
| $\alpha_1, \alpha_2, \beta, \kappa_1, \kappa_2, \lambda, \omega$ | 35246 | 34071 | 34563 | 33606 |
| $\alpha_1, \alpha_2, \beta_1, \beta_2, \kappa_1, \kappa_2, \lambda, \omega$ | 34699 | 33231 | 33931 | 32821 |

\* All *Hybrid* model variants tested used *SARSA* MF model and the *Forward*<sub>1</sub> MS model (see full text for details).
Abbreviations: learning rate for first-stage ( $\alpha_1$ ) and second-stage ( $\alpha_2$ ); $\alpha$ is when $\alpha_1 = \alpha_2$ ; inverse temperature for first-stage ( $\beta_1$ ) and second-stage ( $\beta_2$ ); $\beta$ is when $\beta_1 = \beta_2$ ; perseveration for first-stage ( $\kappa_1$ ) and second-stage ( $\kappa_2$ ); $\kappa$ is when $\kappa_1 = \kappa_2$ ; eligibility trace ( $\lambda$ ); $\omega$ is the model-sensitive weight.
<sup>†</sup> Best fitting *Hybrid* model variant for the respective subject and analysis type.

**Table S6.**
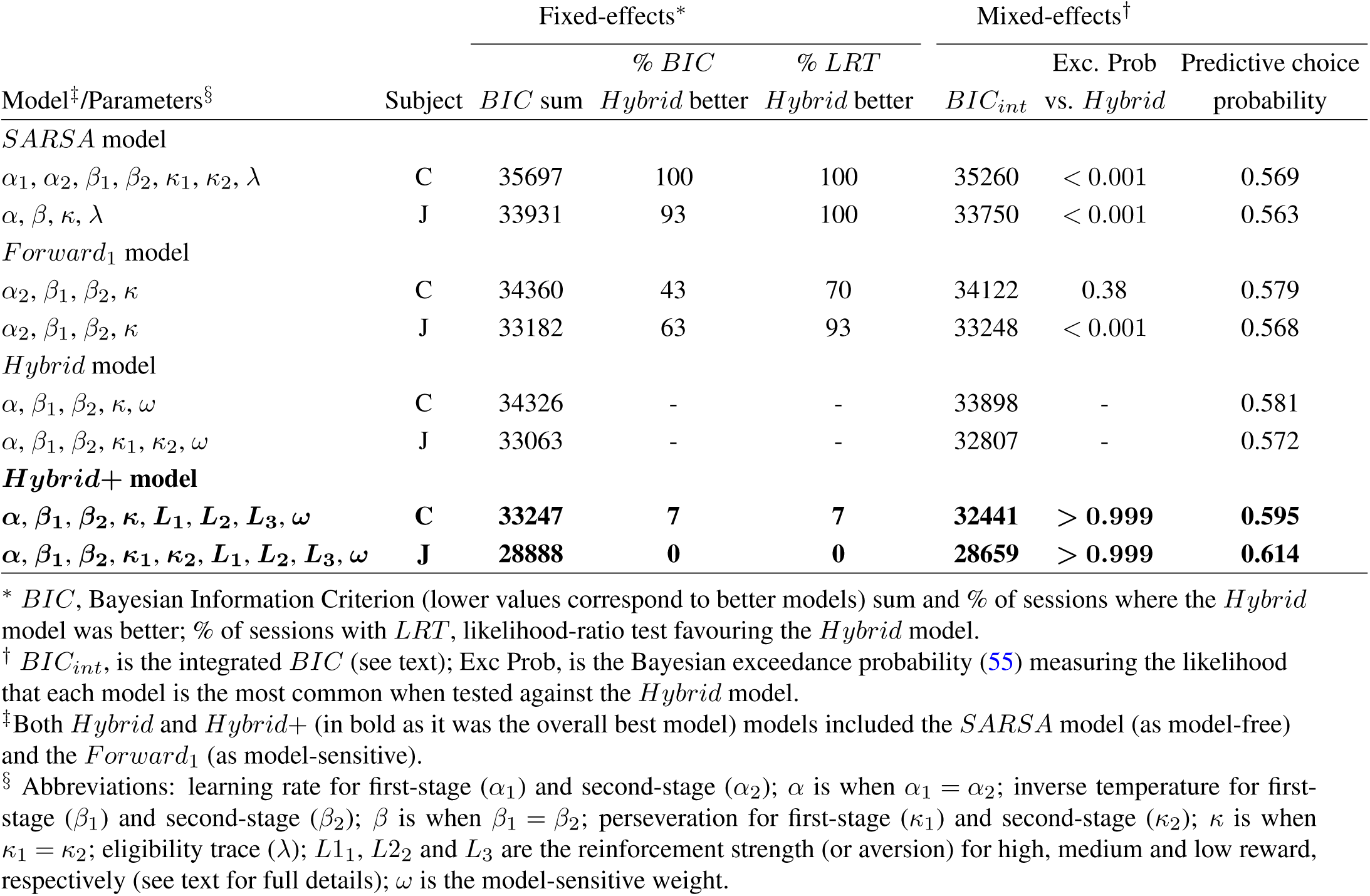
Model comparison results for the three model-sensitive models.

| Model<sup>‡</sup>/Parameters<sup>§</sup> | Subject | Fixed-effects* |  |  | Mixed-effects<sup>†</sup> |  |  |
| --- | --- | --- | --- | --- | --- | --- | --- |
|  |  | *BIC* sum | % *BIC*<br>*Hybrid* better | % *LRT*<br>*Hybrid* better | *BIC*<sub>int</sub> | Exc. Prob<br>vs. *Hybrid* | Predictive choice<br>probability |
| *SARSA* model |  |  |  |  |  |  |  |
| $\alpha_1, \alpha_2, \beta_1, \beta_2, \kappa_1, \kappa_2, \lambda$ | C | 35697 | 100 | 100 | 35260 | < 0.001 | 0.569 |
| $\alpha, \beta, \kappa, \lambda$ | J | 33931 | 93 | 100 | 33750 | < 0.001 | 0.563 |
| *Forward*<sub>1</sub> model |  |  |  |  |  |  |  |
| $\alpha_2, \beta_1, \beta_2, \kappa$ | C | 34360 | 43 | 70 | 34122 | 0.38 | 0.579 |
| $\alpha_2, \beta_1, \beta_2, \kappa$ | J | 33182 | 63 | 93 | 33248 | < 0.001 | 0.568 |
| *Hybrid* model |  |  |  |  |  |  |  |
| $\alpha, \beta_1, \beta_2, \kappa, \omega$ | C | 34326 | - | - | 33898 | - | 0.581 |
| $\alpha, \beta_1, \beta_2, \kappa_1, \kappa_2, \omega$ | J | 33063 | - | - | 32807 | - | 0.572 |
| ***Hybrid+* model** |  |  |  |  |  |  |  |
| $\alpha, \beta_1, \beta_2, \kappa, L_1, L_2, L_3, \omega$ | C | **33247** | **7** | **7** | **32441** | **> 0.999** | **0.595** |
| $\alpha, \beta_1, \beta_2, \kappa_1, \kappa_2, L_1, L_2, L_3, \omega$ | J | **28888** | **0** | **0** | **28659** | **> 0.999** | **0.614** |

\* *BIC*, Bayesian Information Criterion (lower values correspond to better models) sum and % of sessions where the *Hybrid* model was better; % of sessions with *LRT*, likelihood-ratio test favouring the *Hybrid* model.
<sup>†</sup> *BIC*<sub>int</sub>, is the integrated *BIC* (see text); Exc Prob, is the Bayesian exceedance probability (55) measuring the likelihood that each model is the most common when tested against the *Hybrid* model.
<sup>‡</sup> Both *Hybrid* and *Hybrid+* (in bold as it was the overall best model) models included the *SARSA* model (as model-free) and the *Forward*<sub>1</sub> (as model-sensitive).
<sup>§</sup> Abbreviations: learning rate for first-stage ( $\alpha_1$ ) and second-stage ( $\alpha_2$ ); $\alpha$ is when $\alpha_1 = \alpha_2$ ; inverse temperature for first-stage ( $\beta_1$ ) and second-stage ( $\beta_2$ ); $\beta$ is when $\beta_1 = \beta_2$ ; perseveration for first-stage ( $\kappa_1$ ) and second-stage ( $\kappa_2$ ); $\kappa$ is when $\kappa_1 = \kappa_2$ ; eligibility trace ( $\lambda$ ); $L_1, L_2$ and $L_3$ are the reinforcement strength (or aversion) for high, medium and low reward, respectively (see text for full details); $\omega$ is the model-sensitive weight.

**Table S7.** Linear regression results for predictors of first-stage reaction time up to five trials back.

| Predictors<sup>‡</sup> | Fixed-effects* |  | Mixed-effects<sup>†</sup> |  |
| --- | --- | --- | --- | --- |
|  | C | J | C | J |
| Const | 0.02 (< 0.01)<sup>§</sup> | 0.02 (0.01)<sup>§</sup> | 0.02 (0.01)<sup>§</sup> | 0.03 (0.01)<sup>§</sup> |
| $F_t$ | 0.14 (0.04)<sup>§</sup> | 0.41 (0.07)<sup>§</sup> | 0.23 (0.03)<sup>§</sup> | 0.51 (0.05)<sup>§</sup> |
| $R_{t-1}$ | *0.24 (0.04)<sup>§</sup>* | *-0.66 (0.05)<sup>§</sup>* | *0.29 (0.03)<sup>§</sup>* | *-0.69 (0.07)<sup>§</sup>* |
| $T_{t-1}$ | 0.08 (0.02)<sup>§</sup> | 0.03 (0.02) | 0.09 (0.02)<sup>§</sup> | 0.06 (0.02)<sup>§</sup> |
| $R_{t-1} \times T_{t-1}$ | *0.23 (0.05)<sup>§</sup>* | *0.20 (0.03)<sup>§</sup>* | *0.30 (0.04)<sup>§</sup>* | *0.22 (0.04)<sup>§</sup>* |
| $R_{t-2}$ | *-0.14 (0.02)<sup>§</sup>* | *-0.09 (0.02)<sup>§</sup>* | *-0.14 (0.02)<sup>§</sup>* | *-0.04 (0.02)<sup>§</sup>* |
| $T_{t-2}$ | 0.03 (0.02) | -0.03 (0.02) | 0.06 (0.02)<sup>§</sup> | 0.08 (0.01)<sup>§</sup> |
| $R_{t-2} \times T_{t-2}$ | *0.09 (0.04)* | *0.07 (0.03)* | *0.13 (0.03)<sup>§</sup>* | *0.10 (0.03)<sup>§</sup>* |
| $R_{t-3}$ | *-0.07 (0.02)<sup>§</sup>* | *-0.02 (0.02)* | *-0.08 (0.02)<sup>§</sup>* | *-0.03 (0.02)<sup>§</sup>* |
| $T_{t-3}$ | 0.01 (0.02) | -0.01 (0.02) | 0.05 (0.04)<sup>§</sup> | 0.09 (0.02)<sup>§</sup> |
| $R_{t-3} \times T_{t-3}$ | *0.02 (0.04)* | *-0.01 (0.03)* | *0.14 (0.03)<sup>§</sup>* | *0.10 (0.03)<sup>§</sup>* |
| $R_{t-4}$ | *-0.01 (0.02)* | *0.04 (0.02)* | *-0.07 (0.02)<sup>§</sup>* | *-0.04 (0.02)<sup>§</sup>* |
| $T_{t-4}$ | 0.01 (0.02) | 0.01 (0.02) | 0.06 (0.02)<sup>§</sup> | 0.03 (0.01)<sup>§</sup> |
| $R_{t-4} \times T_{t-4}$ | *-0.02 (0.04)* | *0.04 (0.02)* | *0.10 (0.03)<sup>§</sup>* | *0.07 (0.02)<sup>§</sup>* |
| $R_{t-5}$ | *-0.03 (0.02)* | *0.04 (0.02)<sup>¶</sup>* | *0.11 (0.04)<sup>§</sup>* | *-0.04 (0.01)<sup>§</sup>* |
| $T_{t-5}$ | < 0.01 (0.02) | 0.02 (0.02) | 0.06 (0.03) | 0.02 (0.02) |
| $R_{t-5} \times T_{t-5}$ | *0.06 (0.04)* | *0.04 (0.05)<sup>§</sup>* | *0.14 (0.03)<sup>§</sup>* | *0.16 (0.04)* |

\* Values of fixed-effects results are mean (SEM) of the regression coefficients across sessions.
<sup>†</sup> Values of mixed-effects results are the regression coefficients (SE).
<sup>‡</sup> For the given trial $t$ , the variables used were: F was used to model (linearly-increasing) fatigue by counting the trials in the session; C is first-stage choice (1=car picture, 0=watering can picture); R is outcome level (assumed as continuous and with low=1, medium=2, high=3); and T is transition (rare=1, common=0). Const is the constant term. Predictors were mean centred and continuous variables were also scaled by dividing them by two SD (adjustments made before the computation of the interaction terms). In italic are the predictors of interest.
<sup>§</sup> Significance at the 0.01 level.
<sup>¶</sup> Significance at the 0.05 level.

